# Mapping *in silico* genetic networks of the *KMT2D* tumour suppressor gene to uncover novel functional associations and cancer cell vulnerabilities

**DOI:** 10.1101/2024.01.17.575929

**Authors:** Yuka Takemon, Erin D. Pleasance, Alessia Gagliardi, Christopher S. Hughes, Veronika Csizmok, Kathleen Wee, Diane L. Trinh, Ryan D. Huff, Andrew J. Mungall, Richard A. Moore, Eric Chuah, Karen L. Mungall, Eleanor Lewis, Jessica Nelson, Howard J. Lim, Daniel J. Renouf, Steven JM. Jones, Janessa Laskin, Marco A. Marra

## Abstract

Loss-of-function (LOF) alterations in tumour suppressor genes cannot be directly targeted. Approaches characterising gene function and vulnerabilities conferred by such mutations are required. Here, we computationally map genetic networks of *KMT2D*, a tumour suppressor gene frequently mutated in several cancer types. Using *KMT2D* loss-of-function (*KMT2D*^LOF^) mutations as a model, we illustrate the utility of *in silico* genetic networks in uncovering novel functional associations and vulnerabilities in cancer cells with LOF alterations affecting tumour suppressor genes. We revealed genetic interactors with functions in histone modification, metabolism, and immune response, and synthetic lethal (SL) candidates, including some encoding existing therapeutic targets. Analysing patient data from The Cancer Genome Atlas and the Personalized OncoGenomics Project, we showed, for example, elevated immune checkpoint response markers in *KMT2D*^LOF^ cases, possibly supporting *KMT2D*^LOF^ as an immune checkpoint inhibitor biomarker. Our study illustrates how tumour suppressor gene LOF alterations can be exploited to reveal potentially targetable cancer cell vulnerabilities.

## Main

Loss-of-function (LOF) alterations in tumour suppressor genes are conceptually challenging to target therapeutically, due to the reduction or complete loss of the encoded protein that, under normal conditions, provides the cellular restraints necessary to prevent e.g. aberrant proliferation or genomic instability^1^. The functional inadequacy or complete lack of mutated tumour suppressor gene protein renders most inhibitory drugs illogical in contrast to therapies targeting oncogenic proteins^1^. Genetic networks, namely essentiality networks^2–6^ and genetic interaction (GI) networks^7–9^, provide a powerful means to map and interrogate functional interaction landscapes and to identify vulnerabilities of cells harbouring LOF alterations in tumour suppressor genes. Briefly, essentiality networks include co-essential and anti-essential genes, where the perturbation of two genes has similar (synergistic) or opposing (antagonistic) effects on cell survival across various cell lines, respectively^2,4,5^. GI networks include synthetic lethal (SL) and alleviating (AL) interactors, where the perturbation of two genes together in the same cell confers reduced or advantageous fitness effects, respectively, but the perturbation of either gene alone does not affect cell fitness^7^. Genetic network analyses have revealed novel functional relationships between genes, pathways, and cancer cell vulnerabilities^2–6,8,9^. Even so, mapping genetic networks at scale using *in vitro* screening techniques remains a challenge^10,11^.

Here, we report the use of an *in silico* genetic screening approach to systematically characterise tumour suppressor gene function and vulnerabilities of cancer cells harbouring LOF alterations in a tumour suppressor gene. We applied our method to map the genetic networks of *KMT2D,* a frequently mutated tumour suppressor gene across cancer types^12^. Examination of *KMT2D*’s essentiality network revealed novel association with genes that play roles in histone modification, transcription, mitotic cell cycle regulation, glycolysis, and DNA replication, and those encoding proteins that interact with KMT2D on the chromatin. Through mapping cancer type-specific *in silico KMT2D* GI networks, we revealed several SL candidates, namely *WRN*, *MDM2*, *NDUFB4*, and *TUBA1B*, that encode targets of existing and in-development therapeutics, making them potentially viable drug targets in cancers harbouring *KMT2D*^LOF^ alterations. We showed that *KMT2D*^LOF^ cancer cell lines with microsatellite instability (MSI) have aberrant microsatellite expansion, and using The Cancer Genome Atlas (TCGA) data, we showed dysregulated SL candidate associated functions, such as in p53 regulation and metabolism. Furthermore, using both TCGA and whole-genome and transcriptome data from the Personalized OncoGenomics (POG) Program at BC Cancer (NCT021556210^13,14^), a cohort of advanced and metastatic cancer patients that have been heavily treated, we showed that MSI cases with *KMT2D*^LOF^ alterations show significantly elevated immune checkpoint response markers compared to MSI *KMT2D*^WT^ cases, indicating *KMT2D*^LOF^ mutations may be a biomarker for immune checkpoint inhibitor (ICI) treatment stratification. Our work thus models a more general approach, in which *in silico* genetic network maps are used to identify novel functional associations and cancer cell vulnerabilities associated with tumour suppressor gene alterations.

## Results

### KMT2D essentiality network includes genes associated with histone modification, transcription, mitotic cell cycle regulation, glycolysis, and DNA replication

To map *KMT2D*’s essentiality network we used GRETTA, a tool that we developed that leverages the public cancer dependency map (DepMap) data platform to generate genetic network maps^15^. We compared the fitness effects of *KMT2D* knockout (KO) to the effects of knocking out 18,333 genes that were screened by DepMap in 739 cancer cell lines across >30 cancer types (Methods) and identified 1,014 candidate co-essential genes and 940 candidate anti-essential genes in *KMT2D*’s essentiality network (Figure 1A; Supplementary Table S1A). Several co-essential and anti-essential candidates have been previously linked to KMT2D or its functions, and we provide a summary of their KMT2D-associated functions in Table 1. Notably, we did not observe *KMT2C* as a co-essential gene, indicating that KMT2C and KMT2D may be functionally distinct, which supports a previous observation that identified independent functions of these genes^16^. These candidate genes confirm that essentiality networks can capture known protein-protein interactions and functionally related genes.

**Figure 1.**
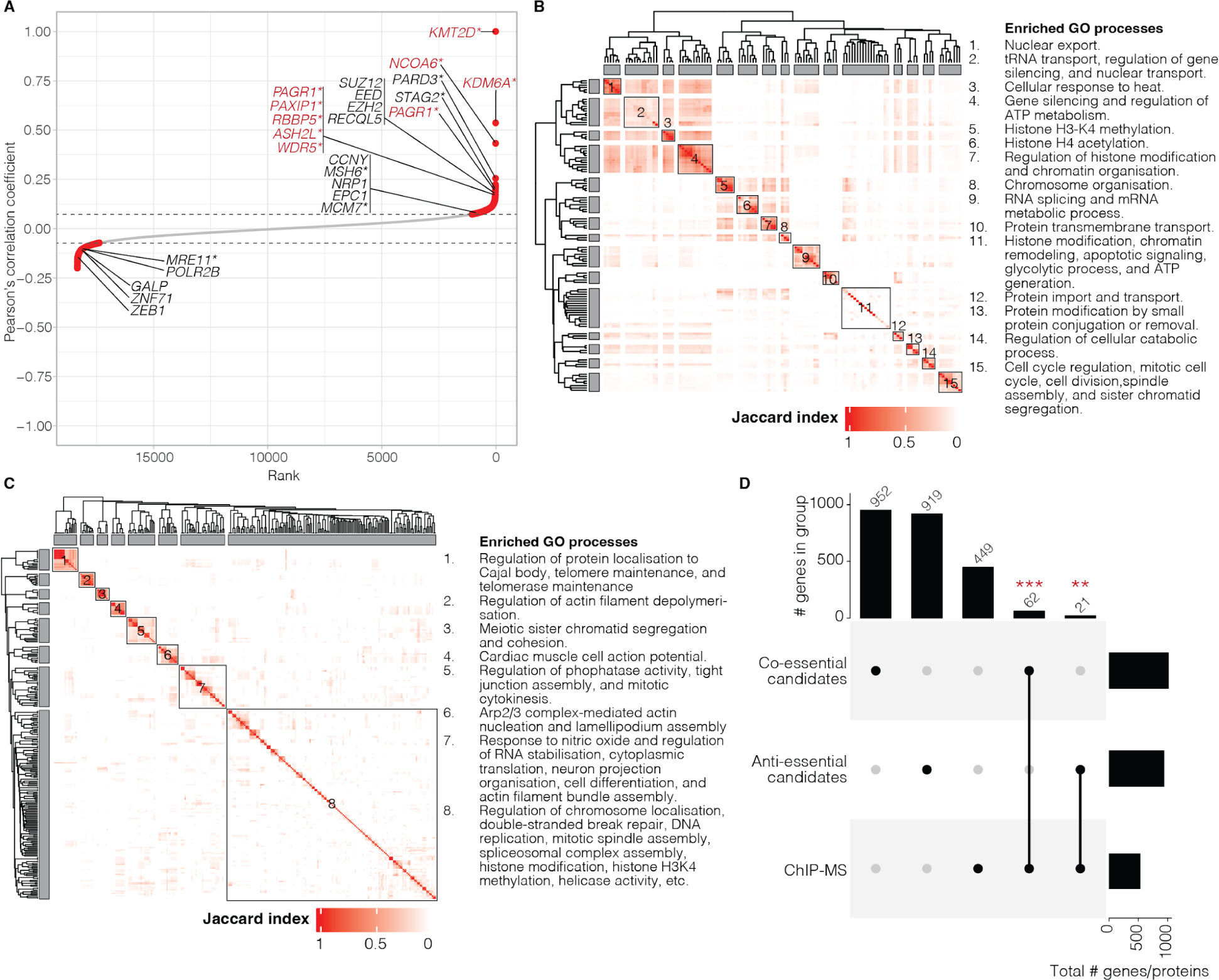
H3K4 methylation, chromatin segregation, and spindle assembly are enriched in *KMT2D* essentiality and protein networks. **A.** Ranked Pearson’s correlation coefficient between *KMT2D* and 18,333 genes. The dotted line denotes the inflection point of the positive and negative curves. Red points indicate candidate co-essential (positive coefficient scores) and anti-essential (negative coefficient scores) genes. Genes labelled in red indicate COMPASS complex members. * indicate KMT2D protein interactors detected in ChIP-MS assay in (C-D). **B-C.** Heatmap showing Jaccard index similarities between 135 significantly enriched GO biological processes from *KMT2D* co-essential candidate genes (B) and 268 significantly enriched GO terms from KMT2D ChIP-MS (C). Jaccard index-based hierarchical clustering was used to group the GO terms into distinct functions (left), and the functions are summarised in the text (right). **D.** Upset plots showing the overlap of genes/proteins detected in KMT2D ChIP-MS and the essentiality network mapping. Bars on the right show the total number of candidates in each row. Bars on the top denote the unique (single black dot) or overlapping (two connected black dots) genes/proteins detected between groups. Fisher’s exact test was to determine significant overlaps: p-value *** < 0.001 and not significant (NS) > 0.05.

**Table 1.** Candidate co-/anti-essential genes and protein interactors with known relations to KMT2D membership/function(s). * indicates genes whose encoded protein were also detected in the KMT2D ChIP-MS analysis.

| Candidates | Candidate membership/function(s) | Known KMT2D membership/function(s) |
| --- | --- | --- |
| *ASH2L* *, *RBBP5* *, *WDR5* *, *KDM6A* *, *NCOA* *, and *PAGR1*, *PAXIP1* * (co-essential) | Core and KMT2D-specific COMPASS protein complex members<sup>17–19</sup>. | COMPASS protein complex member<sup>125</sup>. |
| *CCNY*, *EPC1*, *NRP1*, and *PARD3* * (co-essential)<br>*GALP*, *ZEB1*, and *ZNF71* (anti-essential)<br>KIF5B and RAP1A (ChIP-MS) | Mutations associated with Kabuki and Kabuki-like syndromes<sup>126</sup>. | Loss known to be associated with Kabuki syndrome<sup>127</sup>. |
| *EED*, *EZH2*, and *SUZ12* (co-essential) | Polycomb repressive complex 2 members with roles depositing H3K27me2/3 repressive marks<sup>128</sup>. | Deposits H3K4me3 marks that are known to inhibit PRC2 H3K27me2/3 activity<sup>40,67,129</sup>. |
| *RECQL5* (co-essential) | Known interactor of KMT2D<sup>20</sup>. RecQ DNA helicase with roles in double-stranded DNA break repair, transcription stress, and genomic instability<sup>24,34</sup>. | Known interactor of RECQL5<sup>20</sup>. Associated with transcription stress, and genomic instability<sup>34</sup>. |
| *MRE11* * (anti-essential) | MRN complex member responsible for recognising, repairing, and signalling of double-stranded breaks<sup>130</sup>. | Loss of KMT2D leads to reduced recruitment of MRE11 to replication forks<sup>113</sup>. |
| *POLR2B* (anti-essential)<br>POLR2A (ChIP-MS) | KMT2D interactor encoding RNA polymerase II<sup>20</sup>. | RNA polymerase II interactor. Loss of KMT2D affects POLR2A-associated histone methylation<sup>20</sup>. |
| p53 (ChIP-MS) | Responds to stress stimuli and has mutation-dependent tumour suppressive and oncogenic roles<sup>131</sup>. | p53 and mutant p53 interact with KMT2D<sup>75,76</sup>. |

Next, we performed an enrichment analysis to determine whether there were over-represented biological processes among the candidate co-/anti-essential genes. We identified 135 and eight gene ontology (GO) terms that were significantly enriched in our list of candidate co-essential and anti-essential genes, respectively (q-value < 0.05; Supplementary Table S1B; Methods). We summarised the GO terms enriched in co-essential candidates into 15 distinct groups, based on similarities in genes associated with the terms (Figure 1B; Supplementary Table S1C; Methods). As expected, terms associated with histone modification were enriched. Terms related to cell cycle, cell division, spindle assembly, and sister chromatid segregation were also enriched, including novel candidate co-essential genes and the known *KMT2D*-associated genes *PAGR1*, *PAXIP1^17–19^*, *RECQL5^20^*, *INO80^21^*, and *STAG2^22^*. Notably, these genes also participate in DNA damage repair^23–26^. We also found statistically significantly enriched terms associated with glycolytic processes and ATP generation, which included novel candidate co-essential genes. Among the anti-essential genes, we identified eight significantly enriched GO terms—including DNA-templated transcription initiation, transcription initiation from the RNA polymerase II promoter, histone H3 acetylation, positive regulation of DNA replication, telomere maintenance, telomere organisation, and regulation of telomere maintenance (q-value < 0.05; Supplementary Table S1B; Methods). In addition to KMT2D’s known role in histone modification and transcription initiation^27^, our results indicate that its role may extend into the regulation of cell cycle, cell division, telomere maintenance, glycolysis, and DNA replication.

### KMT2D’s protein interaction network is significantly enriched for functions in histone methylation, chromosome segregation, and DNA replication and repair

To determine whether *KMT2D*’s candidate co-/anti-essential genes also shared physical protein-protein interactions, we performed KMT2D chromatin immunoprecipitation followed by tandem mass spectrometry (ChIP-MS) in human embryonic kidney (HEK293A) cell lines. Given KMT2D’s role as a histone modifier, we chose to capture protein interactions using ChIP to identify chromatin-specific interactions rather than using the common IP-MS techniques, which captures general cellular protein interactions. We identified 532 candidate KMT2D chromatin interactors, which included 14 proteins that were previously described to interact, or have functions that are related to those known to KMT2D (summarised in Table 1; Supplementary Table S2A; Methods).

To highlight functions that were significantly enriched in proteins detected using KMT2D ChIP-MS, we performed an enrichment analysis. 268 GO terms were significantly enriched in candidate KMT2D chromatin interactors (q-value < 0.05; Figure 1C; Methods; Supplementary Table S2B). We summarised these terms into eight distinct groups and identified significant enrichment for proteins associated with the regulation of telomere maintenance, actin filament depolymerisation, sister chromatid segregation and cohesion, DNA replication, and double-stranded break repair (Figure 1C; Supplementary Table S2C; Methods). Additionally, similar to *KMT2D*’s co-essential candidates, protein interactors were also significantly enriched for functions in H3K4 methylation, chromosome segregation, and spindle assembly. Further highlighting their similarities, we found statistically significant overlap between KMT2D protein interactor candidates and co-essential gene candidates (62 proteins/genes overlapped; Fisher’s exact test p-value = 2.18x10^-8^; Figure 1D; Supplementary Table S2D). However, overlap between candidate protein interactors and anti-essential gene candidates were not significant (21 proteins/genes overlapped; Fisher’s exact test p-value = 0.12; Figure 1D). Interestingly, among genes that shared both protein interaction and essentiality with *KMT2D*, we found *OGT* as a co-essential candidate. *OGT* encodes the nutrient-sensing enzyme O-GlcNAc transferase, which has been shown to modify SET1-COMPASS activities in *Drosophila* and interact with SET1A-COMPASS to promote depositions of H3k4me3 marks in mice^28^. OGT has also been implicated in regulation of DNA replication and genomic integrity maintenance^29,30^. However, OGT has not yet been directly associated with KMT2D. Notably, we also found that KMT2D shared protein interaction and gene essentiality with several proteins involved in DNA replication and repair, namely MCM7, MRE11, MSH6, and STAG2. MCM7 and MRE11 have been implicated in DNA replication-coupled repair^31^. MSH6 participates in mismatch repair ^32^. STAG2 is a member of the cohesin complex, responsible for the cohesion of sister chromatids, DNA replication fork progression, and maintaining genome stability^26,33^. Our results thus support the notion that, in addition to its role in histone methylation, KMT2D may also be involved in the regulation of chromosome segregation and DNA replication, which is consistent with previously published evidence showing KMT2D’s role in genome integrity maintenance^20,34^.

### In silico genetic screens reveal pan-cancer and context-specific KMT2D GIs associated with mitotic processes, DNA repair, metabolism, and immune response

We next used GRETTA^15^ to perform pan-cancer and cancer type-specific *in silico* genome-wide KO screens, with the aim of identifying SL and AL interactors of *KMT2D*. To determine the cancer types that were most relevant to *KMT2D*^LOF^ cancers, we queried the datasets from The Cancer Genome Atlas (TCGA; 10,217 tumour samples across 33 cancer types)^35^ and included additional datasets from cancer types that are known to frequently harbour *KMT2D* alterations (see Methods for details). We selected 14 cancer types in which at least 10% of samples harboured *KMT2D* mutations and where 25% of the total *KMT2D* mutations were LOF (Figure 2A; Methods). Next, we selected cancer types with at least ten DepMap cancer cell lines, using the criterion of Behan et al.^8^. This resulted in the identification of nine cancer types (namely bladder urothelial carcinomas [BLCA], B-cell non-Hodgkin lymphoma [B-NHL], colorectal adenocarcinomas [COAD/READ], esophageal carcinomas [ESCA], head and neck squamous cell carcinomas [HNSC], lung squamous cell carcinomas [LUSC], small cell lung cancers [SCLC], stomach adenocarcinomas [STAD], and uterine corpus endometrial carcinoma [UCEC]), in which a cancer type-specific *in silico* GI screen was performed (Figure 2A; Methods). In addition to these nine, we also performed a pan-cancer screen that included all DepMap cell lines. Given that the pan-cancer screen would include cell lines consisting of various cancer types, we expected this screen to contain substantial noise. Also, we expected positive signals (i.e. GI candidates) arising from a pan-cancer screen would represent strong signals, and GI candidates appearing in both pan-cancer and cancer type-specific screens to be high-confidence candidates. Therefore, we rationalised the inclusion of the pan-cancer dataset, which resulted in the analysis of ten datasets.

**Figure 2.**
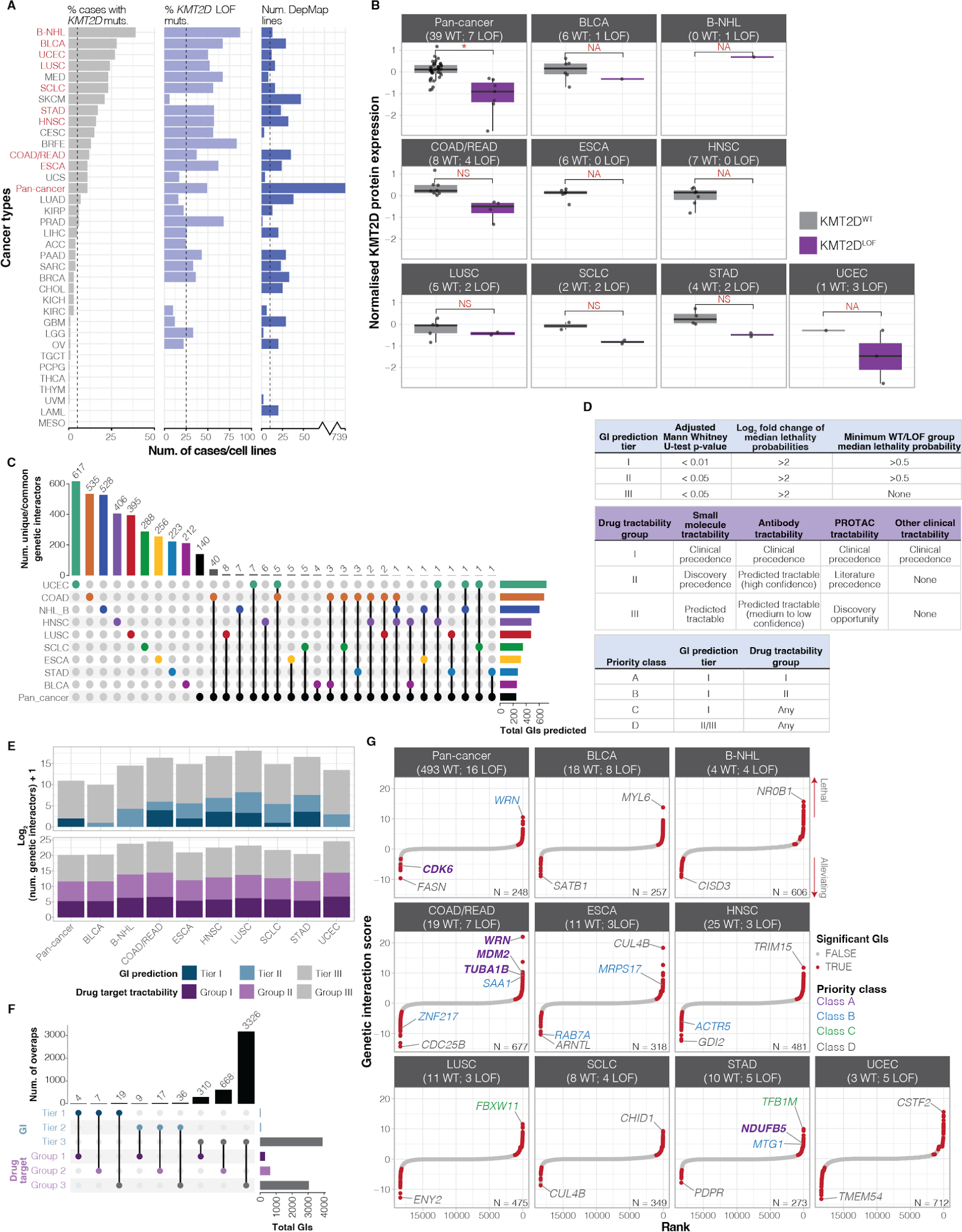
Identifying pan-cancer and cancer type-specific *KMT2D* GI networks. **A.** (Left) Percentage of *KMT2D* small nucleotide mutations (SNVs) detected across 35 cancer types and a pan-cancer dataset. (Middle) Percentage of SNVs that are LOF mutations. (Right) The number of DepMap cancer cell lines for each category. Dotted lines show a threshold for each panel. Red labels indicate the ten datasets that passed all three thresholds and were selected for *in silico* screening. See Methods for cancer type abbreviations. **B.** Normalised KMT2D protein expression for *KMT2D*^WT^ and *KMT2D*^LOF^ DepMap cancer cell lines. **C.** Upset plot showing the number of unique or overlapping candidate GIs between screens. Bars on the right indicate the total number of GIs predicted in each screen. Bars on top indicate the number of GIs that are unique to or shared between the screens. Single coloured dots indicate unique GIs, while linked coloured dots indicate those that are shared between screens. **D.** Definitions for prioritising GI candidates. (Top) GI tiers are based on *in silico* screen statistics. (Middle) Drug groups are based on OpenTarget definitions. (Bottom) Priority classes combine GI tier and drug group classifications, with class A having the highest priority and D having lowest priority candidates. **E.** Bar plots showing the log_2_-transformed number of GI candidates + 1 grouped into GI tiers (top) and drug tractability groups (bottom). **F.** Upset plot showing the number of GI candidates with overlapping priority groups. Right bars indicate the total number of candidates in a GI tier or tractability group. Top bars indicate the number of candidates shared between GI tiers and tractability groups indicated by linked coloured dots. **G.** Ranked GI score summary for each of the ten *in silico* screens. (Bottom right corners) The total numbers of candidates. Class A, B, and the candidate with the highest and lowest GI scores are labelled (see Supplementary Table S3D).

For the pan-cancer dataset and each of the nine cancer type-specific datasets, we used GRETTA^6,15^ to identify *KMT2D*^LOF^ and *KMT2D*^WT^ DepMap cancer cell lines (Supplemental Figure 1A; Supplementary Table S3A; Methods). Next, we compared the *KMT2D* mRNA and protein expression between the *KMT2D*^WT^ and *KMT2D*^LOF^ cell lines within each dataset. *KMT2D* mRNA expression was significantly higher in *KMT2D*^LOF^ lines than *KMT2D*^WT^ lines in the pan-cancer and UCEC datasets (Welch’s t-test p-value < 0.05; Supplemental Figure 1B). On the other hand, KMT2D protein expression in *KMT2D*^LOF^ cell lines was significantly lower in the pan-cancer dataset and trended lower than in *KMT2D*^WT^ cell lines across all nine cancer type-specific datasets for which data were available, including COAD/READ, LUSC, SCLC, STAD, and UCEC (Figure 2B). This indicated that LOF alterations in *KMT2D* were associated with lower protein expression in most cases, as expected, and that the elevated mRNA expression may be a result of a compensatory mechanism induced by deleterious mutations^36,37^. We also compared the relative global levels of active histone marks captured by DepMap, namely H3K4me1, H3K4me2, and H3K27ac, which are known to be reduced in KMT2D-deficient cells^38–42^. Briefly, DepMap utilises a MS-based method to profile relative changes in levels of histone modifications, such as methylation and acetylation, and allows quantification of marks as combinations (e.g. H3K27ac1K36me0), which is not generally possible with antibody-based methods^43^. We found a significant reduction of global H3K4me1 and H3K4me2 levels in *KMT2D*^LOF^ cell lines of the pan-cancer and COAD/READ-specific datasets compared to their respective *KMT2D*^WT^ cell lines (Welch’s t-test p-value < 0.05; Supplemental Figure 1C). B-NHL *KMT2D*^LOF^ cell lines also had significantly reduced levels of H3K4me2 compared to *KMT2D*^WT^ cell lines (Welch’s t-test p-value < 0.05; Supplemental Figure 1C), and both H3K4me1 and H3K4me2 levels trended lower in *KMT2D*^LOF^ BLCA, SCLC, and UCEC cell lines, consistent with a reduction in KMT2D activity. Interestingly, the combined histone mark levels H3K27ac1K36me0 were significantly higher in B-NHL *KMT2D*^LOF^ cell lines compared to *KMT2D*^WT^ cell lines (Welch’s t-test p-value < 0.05; Supplemental Figure 1C), which suggests that KMT2C, EP300, or CREBBP activity may be compensating to maintain homeostasis of transcriptional activity disrupted by KMT2D loss^27^. Altogether, these results are consistent with the notion that KMT2D function is reduced in our selected *KMT2D*^LOF^ cell lines and that such cell lines are suitable for our *in silico* genetic screening analyses.

Finally, using GRETTA^15^, we performed differential lethality analyses to predict *KMT2D* GIs. In this analysis, a gene KO that led to significantly higher lethality probability in the *KMT2D*^LOF^ cell lines compared to *KMT2D*^WT^ cell lines indicated a potential SL interactor, whereas a KO that led to significantly lower lethality probability in *KMT2D*^LOF^ cell lines indicated a potential AL interactor. Using this method, we predicted 4,396 GIs with 3,692 unique GIs across all screens (Supplementary Table S3B; Methods). In context-specific screens, most predicted *KMT2D* GIs (∼80%) were unique to the cancer type in which they were screened (Figure 2C), which is consistent with the view that GIs are highly context-dependent^44^. As expected, 46% of GIs predicted in the pan-cancer screen were also found in context-sensitive screens, given that cell lines in the pan-cancer screens are made up of several cancer types. The largest overlap was seen between the pan-cancer and COAD/READ screens, which is likely due to the large proportion of *KMT2D*^LOF^ cell lines that are COAD/READ (5/16 cell lines).

To highlight pathways that are associated with candidate *KMT2D* GIs, we annotated the SL and AL candidates with the biological processes in which they are involved. The SL candidates were associated with 67 biological processes (Methods; Supplementary Table S3C). Of these, three (4%) were associated with mitotic processes and homologous recombination, five (7%) were associated with metabolic processes, and ten (15%) were associated with T-cell cytotoxicity and immune cell response (Supplementary Table S3C). The AL candidates were associated with 301 biological processes. Five (∼2%) were associated with mitotic processes, 26 (9%) with metabolism, and four (1%) with immune response (Supplementary Table S3C). These results indicate that genes associated with mitotic processes and immune response were more abundant among SL candidates and genes associated with metabolic processes were more abundant in AL candidates. Therefore, *KMT2D*^LOF^ cells may be vulnerable to perturbations in mitotic processes and immune response, whereas they may be more resilient to disruptions in metabolic processes.

### GI prediction scores and target tractability assignments prioritise therapeutically promising KMT2D GIs

Following the method introduced by Behan et al.^8^ to determine high-quality *KMT2D* interactors and prioritise those that might be candidate drug targets, we combined two classification systems, one based on GI prediction statistics and the other based on drug tractability (i.e. the likelihood of identifying a drug targeting the protein encoded by the genetic interactor; summarised in Figure 2D; Methods). We classified GI candidates into three tiers based on statistical significance (GI tiers I, II, and III), where GI tier I candidates represented the most statistically significant group. The second classification was based on drug tractability (drug groups I, II, and III), with group I containing candidates that are targets of existing drugs. Finally, we combined the GI tiers and drug tractability groups to create four priority classes (A, B, C, and D) with priority class A representing the highest level of GI prediction significance and the highest evidence for drug tractability.

We identified three priority class A candidates—namely the SL interactors *MDM2*, *TUBA1B* (COAD/READ), and *NDUFB5* (STAD) and the AL interactor *CDK6* (in the pan-cancer dataset)—that encode targets of approved anticancer drugs or that are the focus of drug development efforts (Figure 2E-G; Supplementary Table S3D)^45^. *MDM2* encodes a negative regulator of p53 that is amplified and overexpressed in 9% of colorectal cancers^46^. *TUBA1B* encodes α-tubulin, a microtubule component that functions in cell motility, adhesion, and cell division. In hepatocellular carcinoma, elevated TUBA1B expression has been associated with a superior response to ICI therapy^47^. In COAD cases, elevated expression was associated with improved overall survival and with CD8+ pre-exhausted T-cells^48^. Interestingly, both MDM2 and TUBA1B have been associated with the DNA damage response^49,50^, a function also associated with KMT2D^20^. *NDUFB5* encodes a subunit of complex I, which plays a role in oxidative phosphorylation^51,52^, a pathway known to be dysregulated in KMT2D-deficient lung adenocarcinomas and Kabuki syndrome patients^53,54^.

Priority class B candidates (targets without drugs in clinical development but with high-confidence evidence supporting target tractability, such as having high-quality ligand structures for pharmaceutical development^45^) include seven genes, specifically the SL interactors *WRN* (pan-cancer and COAD/READ), *MRPS17* (ESCA), and *MTG1* (STAD) and the AL interactors *ZNF217* (COAD/READ), *RAB7A* (ESCA), and *ACTR5* (HNSC; priority class B; Figure 2E-G). *WRN* encodes a DNA helicase and nuclease that functions in DNA replication, repair, and transcription^55–57^ and has recently been identified as a SL vulnerability of MMR-deficient MSI cancers, including some colorectal, endometrial, gastric, and ovarian cancers^8,58–62^. Although *WRN* was categorised as a drug target group II candidate by the Open Target Platform^45^ (Methods), there are currently several drug development programs underway to target WRN^63,64^ in MSI cancer; therefore, we re-classified it as a drug target group I and priority class A gene.

Priority class C candidates consisted of interactors without drugs in clinical development and medium to low-confidence evidence supporting tractability. They included 19 genes, including the SL interactors *FBXW11* (LUSC) and *TFB1M* (STAD; priority class C; Figure 2E-G). Interestingly, FBXW11 is part of an E3 ubiquitin ligase complex involved in ubiquitination and proteasomal degradation in tumorigenesis signalling pathways^65^ and has been shown to co-purify with the polycomb repressive complex 2 (PRC2) members EED^66^, SUZ12, and EZH2^65^, which play an antagonistic role to COMPASS complex members by depositing the repressive histone modification mark H3K27me3^67^. TFB1M is a mitochondrial transcription specificity factor that maintains homeostasis of oxidative phosphorylation and glycolysis^68,69^, pathways that have been shown to be dysregulated in KMT2D-deficient lung cancers^53^, epithelial cells, and Kabuki Syndrome patients^54^. Notably, KMT2D-deficient lung cancer cells depend on proper glycolytic activities for survival^53^, which is compatible with the notion that loss of *TFB1M* may be lethal for KMT2D-deficient cells.

### KMT2D^LOF^ MSI cells require WRN for survival

Given that SL candidates *WRN* in the COAD/READ and pan-cancer screens; *MDM2* and *TUBA1B* in the COAD/READ screen; and *NDUFB5* in the STAD screen were identified as the most promising therapeutic targets (priority class A) for *KMT2D*^LOF^ cancers, we sought to understand the relationship between *KMT2D* and these genes further. Given that *WRN* is a known essential gene in MSI cancer cell lines^8,59–62^, we investigated whether microsatellite status played a role in the SL interaction predicted between *WRN* and *KMT2D*. To this end, we used MSI status to stratify the *WRN* KO lethality probability scores of *KMT2D*^LOF^ cell lines and *KMT2D*^WT^ cell lines from the pan-cancer dataset. We found that lethality probabilities resulting from *WRN* KO were significantly higher in *KMT2D*^LOF^ MSI lines than all three classes of *KMT2D*^WT^ cell lines, including *KMT2D*^WT^ MSI cell lines (ANOVA followed by Tukey’s HSD tests p-value < 0.001; Figure 3A; Methods), indicating that *WRN* KO was lethal in *KMT2D*^LOF^ MSI cell lines but not in *KMT2D*^WT^ MSI cell lines. Furthermore, we found that *KMT2D*^LOF^ MSI lines also had significantly higher *WRN* KO lethality probabilities than *KMT2D*^LOF^ microsatellite stable (MSS) cell lines (ANOVA followed by Tukey’s HSD test p-value < 0.001). This indicated that *WRN* may be required for MSI cancer cell survival only when *KMT2D*^LOF^ mutations were present and that MSI alone may not result in *WRN* dependence, in contrast to initial descriptions^8,59–62^. We also used GRETTA to analyse whether cancer cell lines with *WRN* LOF (*WRN*^LOF^) alterations may require *KMT2D* for survival. We found that *WRN*^LOF^ MSS lines were significantly more susceptible to *KMT2D* perturbation than *WRN* WT (*WRN*^WT^) MSS lines (ANOVA followed by Tukey’s HSD test p-value < 0.01; Figure 3B; Supplementary Table S4; Methods), indicating that cancer cell lines with a *WRN*^LOF^ alteration depend on KMT2D functions for survival and that MSI may not play a role in maintaining viability of *WRN*^LOF^ cell lines. Our results are consistent with the notion that *WRN* is a SL interactor of *KMT2D* and that, in cancer cells, MSI alone may not confer a dependency on WRN for survival.

**Figure 3.**
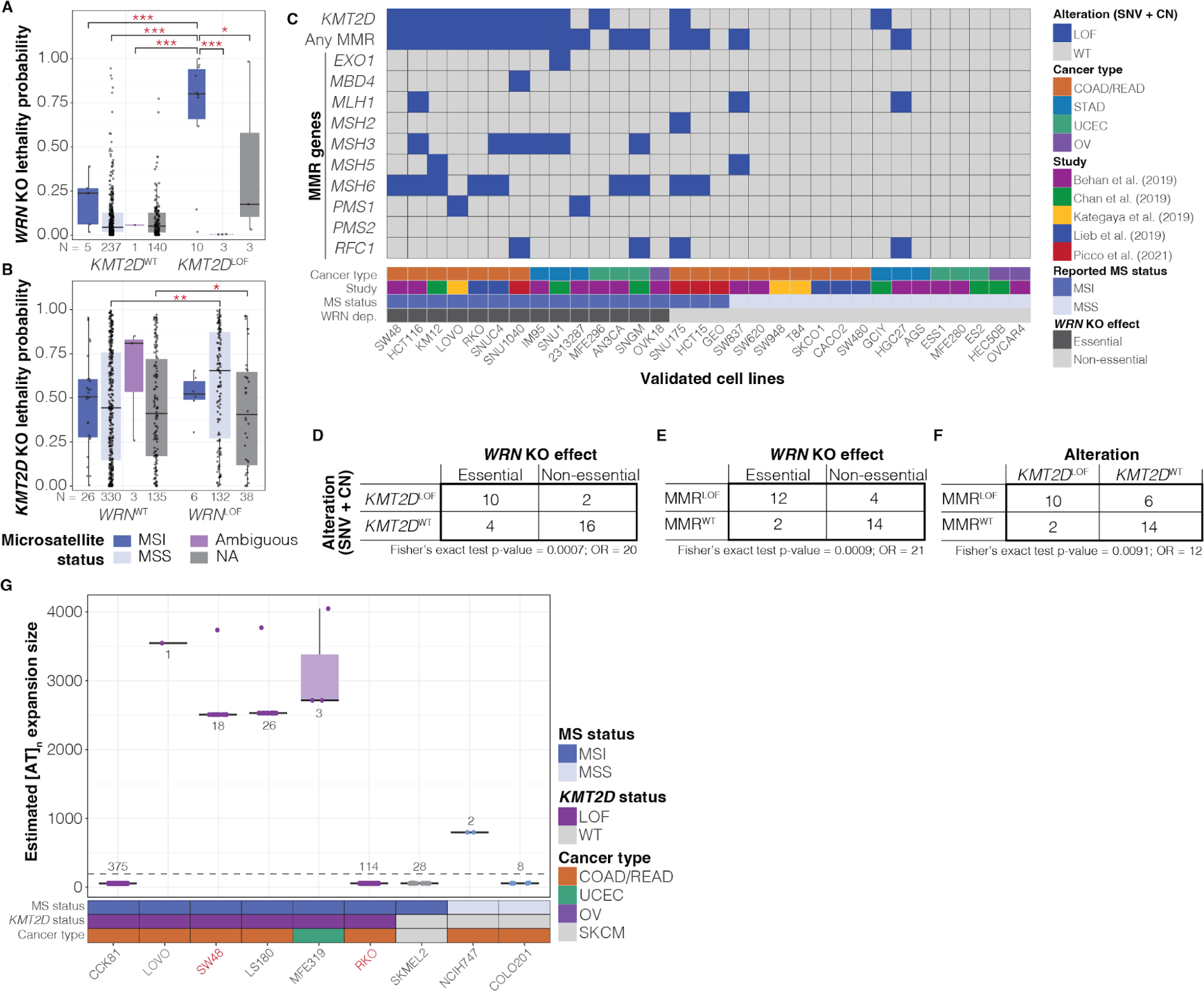
WRN essentiality in MSI cancers is likely mediated by *KMT2D*^LOF^ alterations. A-B. *WRN* (A) and *KMT2D* (B) KO lethality probabilities in *KMT2D* (A) and *WRN* (B) WT and LOF cancer cell lines. Cell lines were separated by microsatellite status as determined in Ghandi et al.^148^. ANOVA followed by Tukey’s HSD p-value * < 0.05, ** < 0.01, *** < 0.001. NA indicates cancer cell lines without MSI/MSS status annotations. **C.** *KMT2D* and MMR gene LOF mutations, detected in cancer cell lines dependent on WRN for viability (see text for references). **D-F.** Contingency tables comparing frequency of LOF alterations in *KMT2D* (D) and MMR genes (E) to *WRN* essentiality and frequency of *KMT2D*^LOF^ mutations to MMR gene mutations (F) in cancer cell lines reported by studies in panel C. **G.** [AT]_n_ microsatellite repeat expansion size estimated by ExpansionHunter Denovo in six DepMap *KMT2D*^LOF^ cell lines from panel A that had WGS data and three *KMT2D*^WT^ lines. Red labels indicate cell lines with *WRN* KO experiments from panel C. Dashed lines indicate the average [AT]_n_ microsatellite repeat expansion size across all events.

Defects in MMR functions as a result of LOF mutations or epigenetic silencing of MMR-associated genes (including *MLH1*, *MLH3*, *MSH2*, *MSH3*, *MSH6*, and *PMS2*) are causes of MSI, resulting in numerous, unrepaired mutations across the genome, mainly in repetitive sequences^70^. Although several studies have shown that WRN is a SL target of MSI cancer cells, restoration of MMR functions through ectopic expression of MMR-associated genes in MSI cell lines only showed partial rescue of cell viability^59^ upon *WRN* KO, and generating MSI conditions through MMR gene KOs in MSS cell lines failed to recapitulate the lethal effects of *WRN* KO^61^. Therefore, we wanted to see whether *KMT2D*^LOF^ alterations in MSI cell lines would lead to a dependency on WRN for survival. To this end, we identified cancer cell lines, in which *WRN* KO was performed and the cell’s viability was validated (i.e. viability assays were performed on cell lines using a single sgRNA targeting *WRN* instead of a pooled sgRNA KO screen), and analysed whether MSI cell lines that harboured *KMT2D*^LOF^ alterations showed more consistent lethal effects upon *WRN* KO compared to *KMT2D*^WT^ MSI cell lines. We identified 32 cancer cell lines (consisting of COAD/READ, STAD, UCEC, and ovarian (OV) cancers) that had genomic sequencing data from DepMap and (in)dependence on WRN for survival validated by studies conducted by either Behan et al.^8^, Chan et al^59^, Kategaya et al.^58^, Lieb et al.^60^, or Picco et al.^62^ (Supplementary Table S5A). For these cell lines, we annotated the presence of LOF alterations in *KMT2D* and MMR genes, namely *EXO1*, *MBD4*, *MLH1*, *MSH2*, *MSH3*, *MSH5*, *MSH6*, *PMS1*, *PSM2*, and *RFC1^62^*, reported MSI status, and the outcomes of *WRN* KO viability assays (Figure 3C; Methods; Supplementary Table S5B). Among the 14 cell lines that showed *WRN* was an essential gene (i.e. that required *WRN* for survival), we identified LOF alterations of MMR genes in 85% (12/14), and concurrent *KMT2D*^LOF^ alterations in 64% (9/14). Among the 18 cell lines where *WRN* was reported as a non-essential gene, we found only 33% with MMR or *KMT2D* alterations (4/18 with MMR gene LOF and 2/18 with *KMT2D*^LOF^). These results indicate that most MMR-deficient lines harbour *KMT2D*^LOF^ alterations and that not all MMR-deficient cell lines require WRN for survival. Furthermore, both *KMT2D*^LOF^ alterations and LOF alteration in MMR genes were significantly enriched in cell lines where *WRN* was essential (Fisher’s exact test p-value = 0.0007 with odds ratio [OR] = 20 and p-value = 0.0009 with OR = 21, respectively; Figure 3D-E). However, LOF alterations in *KMT2D* and MMR also co-occurred (Fisher’s exact test p-values = 0.0091 and OR = 9; Figure 3F), making it difficult to distinguish the independent effects of *KMT2D*^LOF^ alterations and MMR deficiency. Together, these results indicate that a significant number of cell lines that depended on WRN for survival harbour LOF alterations in MMR gene(s) and *KMT2D*.

Next, given that MSI cells require WRN to resolve expanded microsatellite regions containing [AT]_n_ repeat elements for survival and that these expansions could not be recapitulated by MSS cells upon MMR gene KO^61^, we sought to determine whether the loss of *KMT2D* affected the expansion of [AT]_n_ microsatellite regions in MSI cell lines. To this end, we used ExpansionHunter Denovo^71^ (Methods) to profile [AT]_n_ microsatellite regions in WGS data from DepMap cancer cell lines (SRA project: PRJNA523380; Supplementary Table S6A). Briefly, ExpansionHunter Denovo is a catalogue-free method for genome-wide repeat expansion detection and has been used previously to estimate [AT]_n_ repeat element expansion^61^. We characterised microsatellite regions of 14 cell lines, including nine *KMT2D*^LOF^ MSI, two *KMT2D*^WT^ MSI, and two *KMT2D*^WT^ MSS lines, and subsequently filtered for lines that contained more than one anchored in-repeat read (i.e. one read pair maps within and the other outside of a repeat region) in [AT]_n_ regions. This resulted in profiles of nine cell lines, specifically CCK81, LOVO, SW48, LS180, MFE319, RKO, SKMEL2, NCIH747, and COLO201 lines. Interestingly, between *KMT2D*^LOF^ MSI, *KMT2D*^WT^ MSI, and *KMT2D*^WT^ MSS lines, [AT]_n_ sites were estimated to be the most expanded in *KMT2D*^LOF^ MSI cell lines, namely LOVO, SW48, LS180 (COAD/READ), and MFE319 (UCEC; Figure 3G; Supplementary Table S6B). The LOVO^58^ and SW48^8^ lines were previously shown to depend on WRN for survival. Two *KMT2D*^LOF^ MSI lines, CCK81 and RKO (a line that also depends on WRN for survival)^60^, showed below average [AT]_n_ expansion size across events (Figure 3G; mean [AT]_n_ expansion size across samples = 193); however, the number of [AT]_n_ regions detected was among the highest of all samples in both lines. Two *KMT2D*^WT^ lines, SKMEL2 (MSI) and COLO201 (MSS), also showed below-average [AT]_n_ expansion size. The *KMT2D*^WT^ MSS cell line NCIH747 showed above-average [AT]_n_ expansion size; however, the [AT]_n_ size was >50% below that of the expanded *KMT2D*^LOF^ MSI lines. Our results are consistent with the notion that *KMT2D*^LOF^ alteration may exacerbate the genomic destabilising effects of MSI in cancer cell lines and increase the extent or number of expanded [AT]_n_ microsatellite repeat regions, thus contributing to dependency on WRN for cell viability.

### MDM2 is overexpressed in TCGA-COAD/READ KMT2D^LOF^ cases

*MDM2*, or murine double minute 2, encodes an E3 ligase that labels the tumour suppressor p53 for degradation^72^. MDM2 has been shown to be aberrantly upregulated across several cancers, including COAD/READ, leading to enhanced degradation of p53, reduction of p53 activities, and disease progression^72,73^. Although MDM2 has not been linked to KM2TD, MDM2 overexpression has been shown to stabilise and enhance KMT2A accumulation on the chromatin and promote H3K4me3 deposition^74^, and KMT2D has been shown to interact with p53^75,76^. We found a SL interaction between *KMT2D* and *MDM2* in the COAD/READ GI screen and also found a protein-protein interaction between KMT2D and p53. We therefore sought to better understand the relationship between *KMT2D*, *MDM2*, and *TP53*. We first queried TCGA-COAD/READ data to compare the transcriptome profiles of untreated *KMT2D*^LOF^ cases to *KMT2D*^WT^ cases. We found that COAD/READ *KMT2D*^LOF^ cases had significantly higher *MDM2* mRNA expression compared to *KMT2D*^WT^ cases (Welch’s t-test p-value < 0.01; Figure 4A; Supplementary Table S7); however, there was no difference in *TP53* expression (Figure 4B). Interestingly, mutations affecting *KMT2D* and *TP53* were significantly mutually exclusive (Fisher’s exact test p-value < 0.01 and log_2_ odds ratio [OR] -1.25; Figure 4C; Supplementary Table S7), indicating that p53 functions are likely intact in most *KMT2D*^LOF^ cases. Mutual exclusivity between *KMT2D* and *MDM2* was not analysed due to the low number of cases with *MDM2* mutations (N = 7). To determine whether the significantly higher expression of *MDM2* in *KMT2D*^LOF^ cases resulted in reduced p53 function, we compared the expression of genes whose expression is repressed by p53, namely *BIRC5*, *CCNB1*, *CDC25B*, *CDC25C*, *CDKN1A*, *MYC*, *RAD51*, *TERT*, and *VEGFA^77,78^*. *RAD51* expression was significantly elevated (BH-corrected Welch’s t-test p-value < 0.01) and *BIRC5* and *CCNB1* expression showed a trend towards increase in *KMT2D*^LOF^ cases compared to *KMT2D*^WT^ cases (BH-corrected Welch’s t-test p-value < 0.1; Figure 4D). From this, we inferred that p53 function may be reduced in *KMT2D*^LOF^ cases. No difference in expression of *CDC25B*, *CDC25C*, *CDKN1A*, and *MYC* was observed. Conversely, *VEGFA* expression was significantly reduced in *KMT2D*^LOF^ cases compared to *KMT2D*^WT^ cases (BH-corrected Welch’s t-test p-value < 0.1; Figure 4D), which is consistent with previous observations that showed p53 can both repress and promote *VEGFA* expression^79,80^. Altogether, our results show that COAD/READ *KMT2D*^LOF^ cases have increased *MDM2* expression, which may result in reduced or dysregulated p53 tumour suppressor activities, and thus highlight MDM2 as a potentially attractive candidate for therapeutic targeting in COAD/READ *KMT2D*^LOF^ cases.

**Figure 4.**
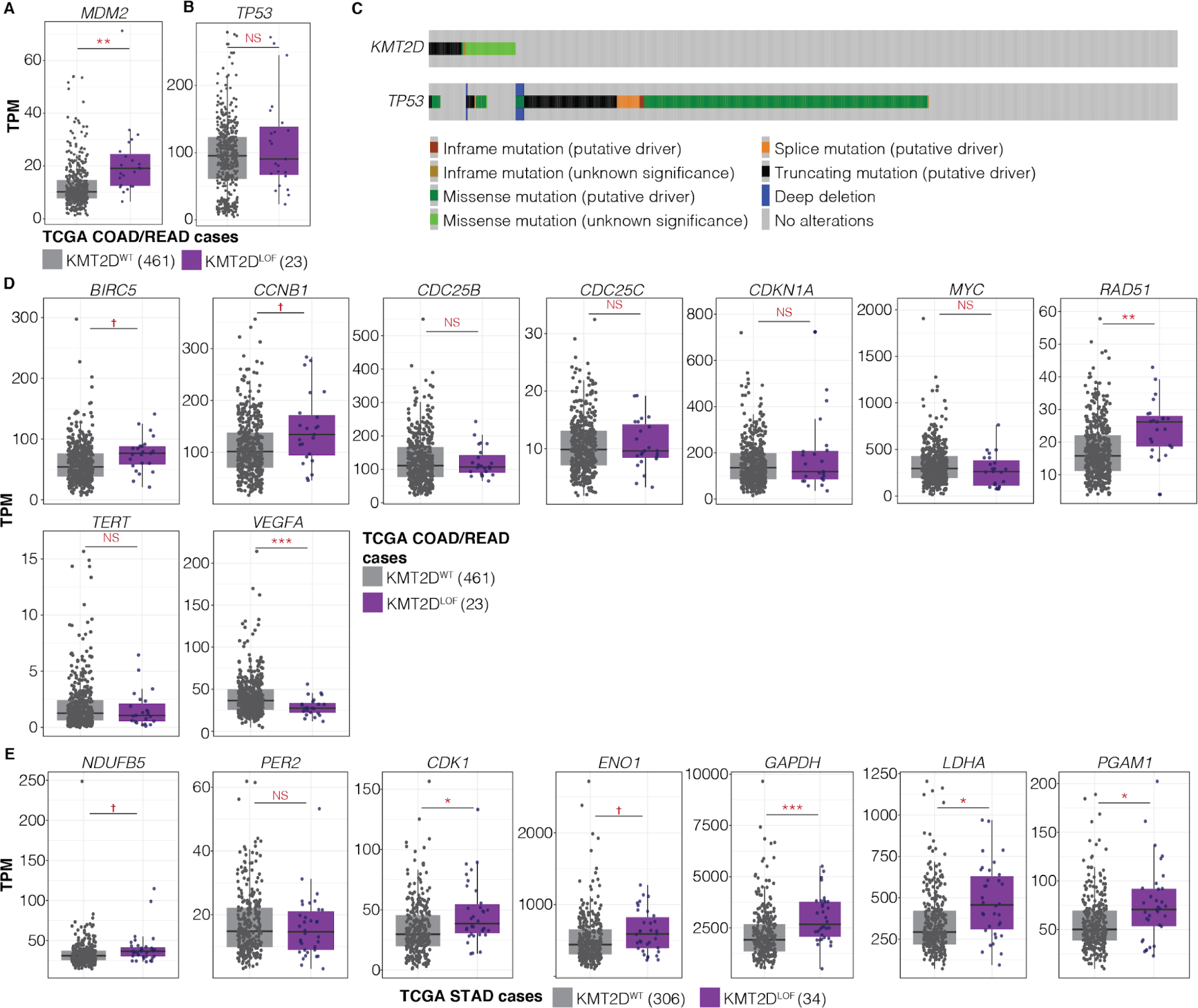
*KMT2D*^LOF^ TCGA cases may be vulnerable to *MDM2* and *NDUFB5* perturbations. **A-B.** *MDM2* (A) and *TP53* (B) mRNA expression in TCGA-COAD/READ *KMT2D*^LOF^ cases and *KMT2D*^WT^ cases. Welch’s t-test p-value ** < 0.01 and NS > 0.05. **C.** OncoPrint of *KMT2D* and *TP53* mutations found in 565 TCGA-COAD/READ cases^204,205^. Fisher’s exact test p-value < 0.01 and log_2_OR = -1.25. **D-E.** (D) mRNA expression of genes regulated by p53 in TCGA-COAD/READ cases and (E) *NDUFB5* and glycolytic genes in TCGA-STAD cases. BH-corrected Welch’s t-test p-value ✝ < 0.1, ** < 0.01, *** < 0.001, and NS > 0.1.

### Glycolytic genes are dysregulated in TCGA-STAD KMT2D^LOF^ cases

*NDUFB5* encodes a subunit of the mitochondrial respiratory chain enzyme Complex I (NADH:ubiquinone oxidoreductase), which functions to create a proton gradient for ATP synthesis through oxidative phosphorylation (OXPHOS)^51,52^. Interestingly, studies by Alam et al.^53^ in a lung cancer model and Maitituoheti et al. ^81^ in a melanoma model showed that KMT2D deficiency led to dysregulation of OXPHOS and glycolytic activity, conferring a vulnerability to perturbation in glycolytic functions. Alam et al.^53^ also demonstrated that KMT2D loss impaired epigenetic signals, which reduced PER2 expression, a transcriptional repressor of multiple glycolytic genes. Therefore, we investigated whether *KMT2D*^LOF^ alterations were associated with differences in expression of *NDUFB5*, *PER2*, and PER2-regulated glycolytic genes in TCGA-STAD cases. We found a trend towards increased *NDUFB5* mRNA expression (BH-corrected Welch’s t-test p-value < 0.1; Figure 4E; Supplementary Table S8; Methods) in *KMT2D*^LOF^ cases compared to *KMT2D*^WT^ cases. Although we did not find a difference in *PER2* expression, PER2-regulated glycolytic genes, namely *CDK1*, *ENO1*, *GAPDH*, *LDHA*, and *PGAM1^53^*, all showed either a trend towards or a statistically significant mRNA expression elevation in *KMT2D*^LOF^ cases compared to *KMT2D*^WT^ cases, consistent with results demonstrated by Alam et al.^53^ in a *Kmt2d* knockdown mouse lung cancer cell line (Figure 4E; Supplementary Table S8; Methods). Furthermore, we found that, in addition to *NDUFB5*, all GI tier I candidates in the STAD screen, namely *ATP2B1^82^*, *DARS2^83^*, *MTG1^84^*, *NUBPL^85^*, and *TFB1M^68,69,86^*(Supplemental Table S3), were SL interactors and were associated with mitochondrial and metabolic functions. Our results further support the notion that *KMT2D* may play a role in metabolism and that *KMT2D*^LOF^ alterations in STAD cases may also confer a vulnerability to OXPHOS or glycolytic perturbations.

### KMT2D^LOF^ MSI TCGA-COAD/READ cases have significantly elevated ICI response indicators

MSI is an approved tissue/site-agnostic biomarker for ICI treatments^87,88^. However, a recently completed analysis of KEYNOTE-177 (in April of 2022), a randomised, open-label, phase-three study comparing MSI/MMR-deficient metastatic colorectal cancer cases treated with pembrolizumab versus chemotherapy, showed that although pembrolizumab-treated patients had fewer treatment-related adverse effects and a progression-free survival improvement was noted, there was no significant difference in overall survival between the two treatment groups^89^. Therefore, there is a need to improve ICI treatment stratification. Given that we identified several SL candidates with roles in T-cell cytotoxicity and the immune response (Supplementary Table S3C), identified *TUBA1B* as a priority class A target in the COAD/READ screen, and showed that MSI can elicit specific vulnerabilities in *KMT2D*^LOF^ cancer cell lines, we sought to understand the impact of *KMT2D*^LOF^ on immune markers in MSS and MSI COAD/READ cases and whether *KMT2D* mutational status might be relevant in the context of ICI treatment. To this end, we profiled the clinical, mutational, and transcriptomic profiles of untreated COAD/READ MSI cases from TCGA. We identified 43 MSI COAD/READ cases (27 *KMT2D*^WT^ MSI and 16 *KMT2D*^LOF^ MSI cases; Methods) and observed a trend towards lower overall survival probability in *KMT2D*^LOF^ MSI cases compared to *KMT2D*^WT^ MSI cases (log-rank test p-value = 0.10; Figure 5A; Supplementary Table S9; Methods). A survival analysis was not performed for MSS cases due to the small sample size of *KMT2D*^LOF^ cases (417 *KMT2D*^WT^ MSS and two *KMT2D*^LOF^ MSS cases). This supports previous observations showing that KMT2D deficiency leads to inferior survival outcomes compared to KMT2D-proficient patients with lymphoma, lung, and breast cancers^90–93^.

**Figure 5.**
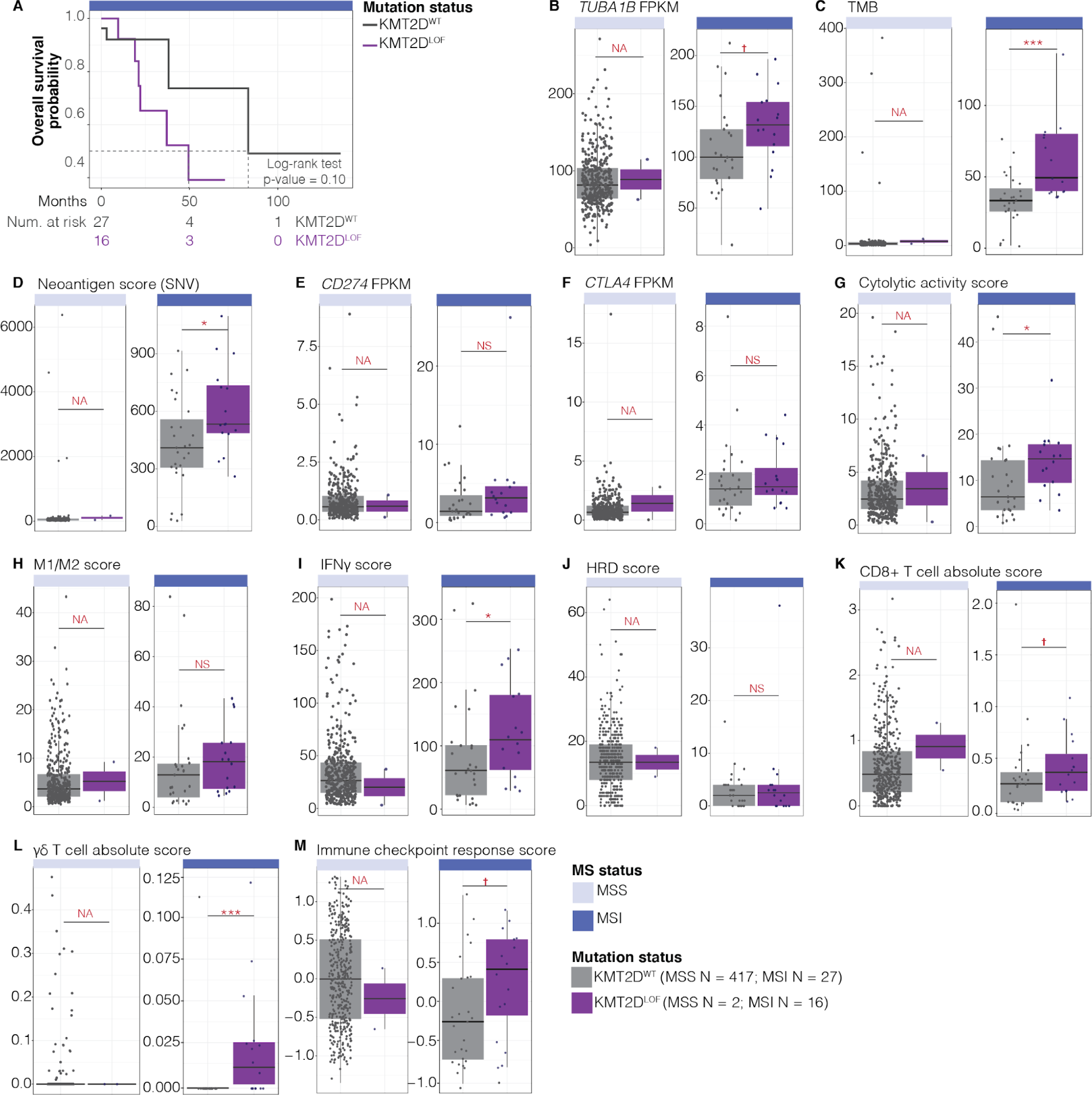
*KMT2D*^LOF^ TCGA-COAD/READ cases may be vulnerable to ICI treatment. **A.** Kaplan–Meier curves (top) and risk table (bottom) of overall survival of MSI TCGA-COAD/READ *KMT2D*^WT^ and *KMT2D*^LOF^ cases. **B-M.** A comparison of ICI response markers between TCGA-COAD/READ *KMT2D*^WT^ and *KMT2D*^LOF^ MSI/MSS cases. ICI response markers included: (B) *TUBA1B* mRNA expression, (C) TMB, (D) neoantigen score, (E-F) mRNA expression of *CD274* and *CTLA4*, (G) cytolytic activity score, (H) M1/M2 macrophage score, (I) IFN*γ* score, (J) HRD score, (K-L) CD8 + and γδ T-cell infiltration score, and (M) PredictIO ICI response score. Welch’s t-test p-value ✝ < 0.1, * < 0.05, ** < 0.01, *** < 0.001, and NS > 0.1. MSS cases were not analysed statistically (NA) due to the small number of *KMT2D*^LOF^ cases.

Next, in the TCGA-COAD/READ cohort, we characterised and compared the expression of *TUBA1B*, as elevated expression has been associated with superior response to ICI therapy^47^, and markers of immune response, including TMB^94,95^, neoantigen scores^96^, *CD274/PD-L1* and *CTLA4* expression^97^, cytolytic activity scores^98,99^, M1/M2 macrophage scores^100,101^, interferon-gamma (IFN*γ*) signalling scores^102^, and homologous recombination deficiency (HRD) scores^103^. Due to a small *KMT2D*^LOF^ MSS sample size, statistical analyses of MSS cases were not performed for any of the immune markers. However, in MSI cases, we found a trend towards higher *TUBA1B* expression in *KMT2D*^LOF^ MSI cases compared to *KMT2D*^WT^ MSI cases (Welch’s t-test p-value = 0.057; Figure 5B; Supplementary Table S9; Methods). We also observed a significantly higher TMB in *KMT2D*^LOF^ MSI compared to *KMT2D*^WT^ MSI cases (Welch’s t-test p-value < 0.001; Figure 5C; Supplementary Table S9; Methods), which supports previous observations showing that KMT2D deficiency led to increased genomic instability in mouse embryonic fibroblast and human colorectal cell line models^20^. A comparison of mutational signatures using the Catalogue Of Somatic Mutations In Cancer (COSMIC) indicated that the single base pair substitution (SBS) signatures associated with MMR deficiency were not significantly different between *KMT2D*^LOF^ and *KMT2D*^WT^ MSI cases (BH-corrected Welch’s p-value > 0.05; Supplementary Table S10; Methods). However, several other mutational signatures, namely SBS1, SBS5, SBS22, and SBS54, were elevated in *KMT2D*^LOF^ MSI cases compared to *KMT2D*^WT^ MSI cases (uncorrected Welch’s p-value < 0.05 but > 0.05 when using BH correction; Supplemental Figure 2A). SBS54 is a possible sequencing artefact, whereas SBS22 is associated with aristolochic acid exposure^104,105^. Interestingly, increased SBS1 and SBS5 mutation rates are associated with increased mutational burden and have been linked to clock-like signatures^104,105^. However, a comparison of chronological age did not show significant difference between *KMT2D*^LOF^ and *KMT2D*^WT^ cases (Welch’s t-test p-value > 0.05; Supplemental Figure 2B). SBS1 arises from an endogenous mutational process that generates guanine:thymine mismatches in double-stranded DNA due to the deamination of 5-methylcytosine to thymine^104–106^. Therefore, the elevated SBS1 signature in *KMT2D*^LOF^ cases indicates that LOF alterations in *KMT2D* may induce endogenous mismatches in addition to those acquired due to MSI. We also observed significantly higher neoantigen scores in *KMT2D*^LOF^ MSI cases than in *KMT2D*^WT^ MSI cases in COAD/READ (Welch’s t-test p-value < 0.05; Figure 5D; Supplementary Table S9; Methods). Next, we compared the expression of *CD274* (encoding PD-L1) and *CTLA4* and observed a trend towards higher *CD274* expression (Welch’s t-test p-value = 0.15; Figure 5E) but no difference in *CTLA4* expression (Welch’s t-test p-value = 0.45; Figure 5F; Supplementary Table S9; Methods) in *KMT2D*^LOF^ MSI cases compared to *KMT2D*^WT^ MSI cases. Furthermore, we also compared the difference in immune cytolytic activity scores^98,99^ (i.e. a measure for anti-tumour immune cell activity) and found that *KMT2D*^LOF^ MSI cases had significantly higher immune cytolytic activity scores compared to *KMT2D*^WT^ MSI cases (Welch’s t-test p-value < 0.05; Figure 5G; Supplementary Table S9; Methods). Together with the trending increase in *CD274* expression, this may indicate that *KMT2D*^LOF^ MSI COAD/READ cancers have increased T-cell infiltration and anti-tumour immune cell activity compared to *KMT2D*^WT^ MSI cases. We next compared M1/M2 macrophage scores. While we did not find a statistically significant difference, M1/M2 scores did show elevated trends in *KMT2D*^LOF^ MSI COAD/READ cases compared to *KMT2D*^WT^ MSI COAD/READ cases (Figure 5H; Supplementary Table S9; Methods). We calculated the IFN*γ* signalling scores^107^ and found that *KMT2D*^LOF^ MSI cases had significantly higher scores than *KMT2D*^WT^ MSI (Welch’s t-test p-value < 0.05; Figure 5I; Supplementary Table S9; Methods). We also compared HRD scores and did not observe a significant difference in HRD scores between *KMT2D*^LOF^ MSI and *KMT2D*^WT^ MSI cases (Figure 5J; Supplementary Table S9; Methods).

Given that tumour immune microenvironment composition has also been shown to be predictive of ICI response^108^, we used CIBERSORTx ^109^ to estimate the extent of immune cell infiltration in TCGA COAD/READ cases and compared the difference in predicted immune cell composition between *KMT2D*^LOF^ MSI and *KMT2D*^WT^ MSI cases. We found a higher predicted abundance of cytotoxic CD8+ T-cells and γδT-cells, a cytotoxic effector that produces pro-inflammatory cytokines^110,111^, in *KMT2D*^LOF^ MSI compared to *KMT2D*^WT^ MSI COAD/READ cases (Welch’s t-test p-value < 0.1 and p < 0.001, respectively; Figure 5K-L; Supplementary Table S9; Methods). However, we did not observe differences in the proportion of other immune cell populations (Supplemental Figure 2C). These results support the notion that MSI COAD/READ cases harbouring *KMT2D*^LOF^ alterations may have more T-cell infiltration than *KMT2D*^WT^ cases.

Finally, to predict whether *KMT2D*^LOF^ MSI COAD/READ cases might respond more favourably to ICI treatment than *KMT2D*^WT^ MSI COAD/READ cases, we calculated their ICI response scores using PredictIO^112^. Briefly, PredictIO uses a signature based on 100 genes that best predict response to ICIs across various cancer types^112^. While we did not find a statistically significant difference, we did observe a higher trend in ICI response scores in *KMT2D*^LOF^ MSI cases (Welch’s t-test p-value < 0.1; Figure 5M; Supplementary Table S9; Methods), indicating that *KMT2D*^LOF^ MSI COAD/READ cases may exhibit a more favourable response when treated with ICIs than *KMT2D*^WT^ MSI cases. Our results thus showed, for the first time, that *KMT2D*^LOF^ alterations in MSI cases appear to confer significantly higher TMB than MSI alone. Furthermore, we showed that *KMT2D*^LOF^ MSI cases had significantly higher expression of neoantigens, immune-activation signatures, cytotoxic T-cell proportions, and ICI-response scores, all consistent with the possibility of more favourable ICI responses compared to *KMT2D*^WT^ MSI COAD/READ cases.

### MSI KMT2D^LOF^ alterations are also associated with elevated trends in immune response indicators in the POG cohort

We next analysed a cohort of advanced and metastatic cancer patients from the Personalized OncoGenomics (POG) Project at BC Cancer (NCT02155621^13,14^). POG cases differ from untreated TCGA cases, as they are at an advanced stage, are incurable, and have been heavily treated. They closely represent the patient populations typically receiving second- or third-line immunotherapy. We first identified COAD/READ cases harbouring *KMT2D*^LOF^ mutations and their MS status. However, the number of MSS and MSI cases harbouring *KMT2D*^LOF^ alterations was small (six and one case, respectively; Figure 6A). Therefore, we also included all solid cancers, which included COAD/READ and 26 other cancer types (Figure 6A-B), and identified 137 *KMT2D*^LOF^ MSS cases and five *KMT2D*^LOF^ MSI cases (Figure 6A). Similar to the trends seen in TCGA-COAD/READ MSI cohort, we computed statistically significantly lower overall survival probability in POG *KMT2D*^LOF^ MSS solid cancer cases than in *KMT2D*^WT^ MSS solid cancer cases (Log-rank test p-value = 0.01; Figure 6C; Supplementary Table S11). However, we did not conduct survival analyses in COAD/READ MSS and MSI solid cancer cases, due to limited case numbers. Our analysis thus showed that the heavily treated POG patients have a similar reduction of overall survival probabilities in patients with *KMT2D*^LOF^ compared to *KMT2D*^WT^ alterations, as seen in untreated TCGA cases.

**Figure 6.**
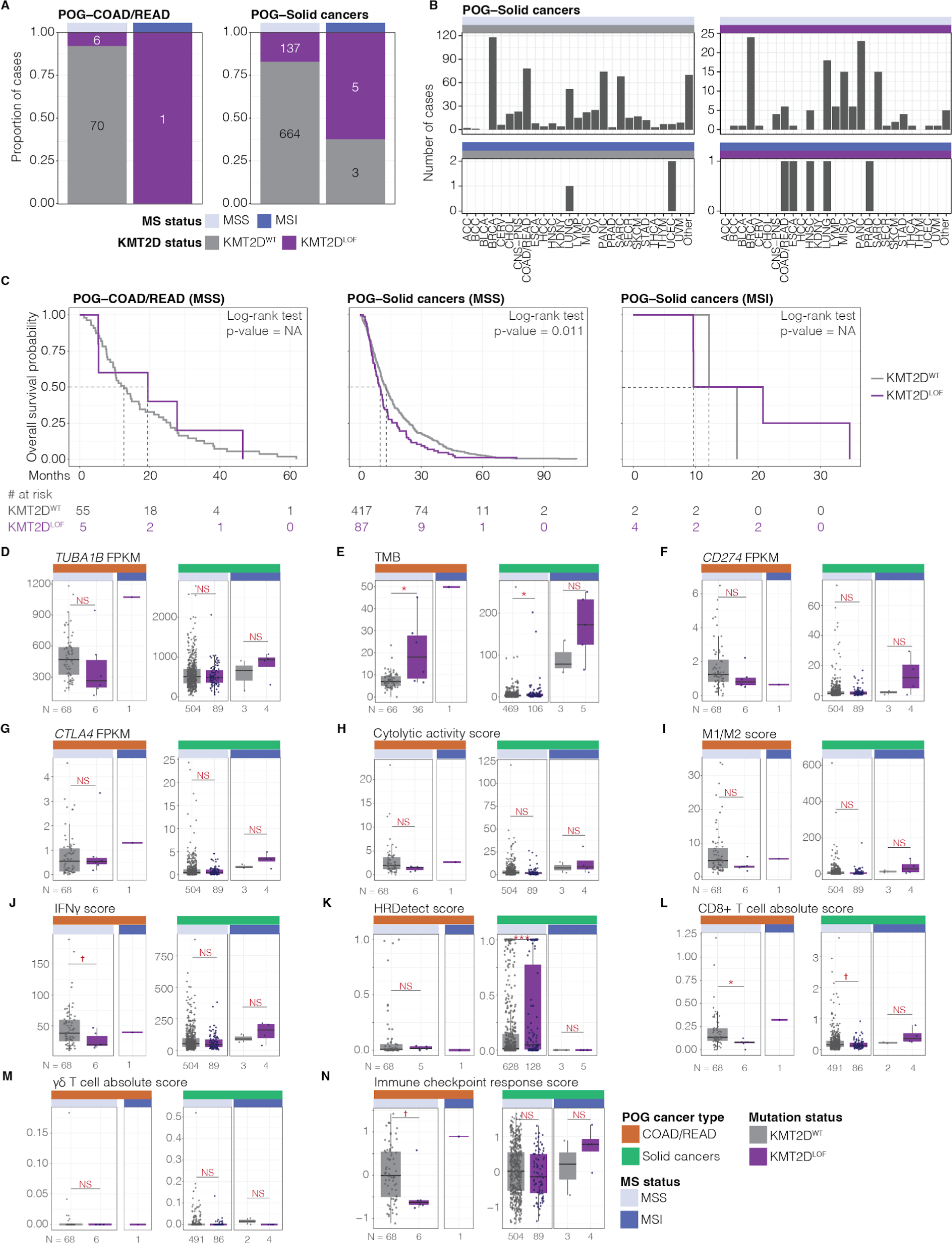
ICI response markers are elevated in *KMT2D*^LOF^ POG-MSI solid cancer cohort cases. **A.** Proportion of advanced and metastatic POG-COAD/READ (left) and all solid cancer (right) cases with MSI and *KMT2D*^LOF^ alterations. **B.** Number of cases by cancer type found in the POG-solid cancer cohort in panel A. **C.** Kaplan–Meier curves (top) and risk table (bottom) comparing overall survival of *KMT2D*^LOF^ cases to *KMT2D*^WT^ cases. **D-N.** A comparison of ICI response markers between *KMT2D*^WT^ and *KMT2D*^LOF^ MSI/MSS cases in POG-COAD/READ and solid cancer cohorts. ICI response markers included: (D) *TUBA1B* mRNA expression, (E) TMB, (F-G) RNA expression of *CD274* and *CTLA4*, (H) cytolytic activity score, (I) M1/M2 macrophage score, (J) IFN*γ* score, (K) HRDetect score (L-M) CD8 + and γδ T-cell infiltration score, and (N) PredictIO ICI response score. Welch’s t-test p-value ✝ < 0.1, * < 0.05, ** < 0.01, *** < 0.001, and NS > 0.1. No statistical analysis was performed for the MSI POG-COAD/READ case, due to limited sample size (N < 3).

Next, we analysed ICI response markers in POG cases (Figure 6D-N; Supplemental Figure 3A-C; Supplementary Table S11; Methods), as we did with TCGA cases. No comparisons were made in COAD/READ MSI cases for any ICI response markers since there was only one case harbouring *KMT2D*^LOF^ alterations, as mentioned above. In COAD/READ and solid cancers, we saw significantly higher TMB in *KMT2D*^LOF^ MSS cases compared to *KMT2D*^WT^ MSS cases (Welch’s t-test p-value < 0.05) and, in solid cancer MSI cases, *KMT2D*^LOF^ cases trended towards elevated TMB compared to *KMT2D*^WT^ cases (Figure 6E). We also saw trends in the same direction for *TUBA1B*, *CD274* and *CTLA4* expression (Figure 6D, F-G); M1/M2 scores (Figure 6I); IFN*γ* scores (Figure 6J); CD8+ T-cell scores (Figure 6L); and ICI response scores (Figure 6N) in *KMT2D*^LOF^ MSI solid cancer compared to *KMT2D*^WT^ MSI solid cancer cases. The substantially smaller sample sizes (a total of one *KMT2D*^LOF^ case in the COAD/READ MSI cases and five in the solid cancer MSI cases) resulted in insufficient power to detect significance differences in the POG cohort. Even so, our findings are consistent with a similar effect of *KMT2D*^LOF^ in the POG cohorts as we observed in TCGA. Our results reinforce the notion that MSI cases with *KMT2D*^LOF^ alterations may respond more favourably to ICI treatment than MSI *KMT2D*^WT^ cases, and that consideration of *KMT2D* mutational status may improve patient stratification for ICI treatment.

## Discussion

In this study, we demonstrated a framework for revealing tumour suppressor gene functions and vulnerabilities of cancer cells harbouring LOF alterations in them. Using the tumour suppressor gene *KMT2D* as an example for our framework, we performed *in silico* genetic network analyses to gain further insights into its functions and identify its cancer type-specific genetic interactors. We identified genes in *KMT2D*’s essentiality network and proteins in its proteomic network that were known interactors and also new candidate interactors that have roles associated with regulation of cell cycle, cell division, telomere maintenance, chromosome segregation, DNA replication and metabolism, thus expanding the extent of KMT2D’s role in these processes. We identified SL and AL candidates that have roles in mitotic processes, metabolic processes, and immune response. Furthermore, results from the *in silico* GI screens identified several SL candidates—namely *WRN* (in the pan-cancer and COAD/READ screens), *MDM2* and *TUBA1B* (in the COAD/READ screen), and *NDUFB5* (in the STAD screen)—which encode proteins that can be inhibited using existing and in-development drugs. We also showed that KMT2D may mediate MSI cell line dependency on WRN for survival and that KMT2D dysregulation may be needed to expand [TA]_n_ microsatellite regions. Using TCGA-COAD/READ cases, we showed that the mRNA expression of *MDM2*, a p53 inhibitor, was significantly increased and the expression of p53 downstream targets were dysregulated in *KMT2D*^LOF^ cases. Using TCGA-STAD cases, we showed that *NDUFB5* expression was elevated and glycolytic genes were significantly elevated in *KMT2D*^LOF^ cases. Interestingly, in addition to showing genes associated with immune response as *KMT2D* SL interactors, which included *TUBA1B*, we also showed that ICI response makers were elevated in *KMT2D*^LOF^ cases. Using untreated TCGA-COAD/READ cases, we showed that MSI *KMT2D*^LOF^ cases have a trending increase in *TUBA1B, CD274*, and *CTLA4* mRNA expression, CD8+ T-cell infiltration scores, and favourable ICI treatment response scores and have significantly higher TMB, neoantigen scores, cytolytic activity, IFN*γ* signalling scores, and γδ T-cell infiltration scores compared to *KMT2D*^WT^ cases. In our retrospective analysis of heavily treated POG cohorts, we found similar trends as in TCGA cohorts. Our study expands KMT2D’s role in genomic stability, metabolism, and immune response; presents promising SL interactors that can potentially be targeted using approved and in-development drugs; and proposes ICI as a possible therapeutic avenue for *KMT2D*^LOF^ cases.

Although KMT2D is best characterised for its role as a histone methyltransferase, loss of *KMT2D* has been associated with increased genomic instability, dysregulation of DNA repair/replication^20,113^, and metabolic dysregulation^53,54^. However, the extent of KMT2D’s involvement in regulating genomic stability and metabolism is still unclear. In addition to supporting KMT2D’s known role as a histone modifier, our analysis of *KMT2D*’s co-essential network and chromatin-specific proteome interaction network reveal an enrichment of genes/proteins related to cell cycle processes, mitotic segregation, and DNA replication/repair that have not previously been linked to KMT2D. These novel candidate SL interactors were also involved in mitotic regulation and homologous recombination. These processes are known to affect genomic stability^114^, and thus further implicating KMT2D in this role.

For the first time, we predicted *KMT2D* candidate SL interactors, namely *WRN*, *MDM2*, *NDUFB5*, and *TUBA1B*, that encode targets of existing drugs or those that are in development. The SL interaction between *KMT2D* and *WRN* contributes a new layer of understanding in MSI cancer vulnerability. Several studies have previously implicated *WRN* to be an essential gene for MSI cancer cell lines^8,59–62^; however, efforts to restore and induce *WRN* dependency were problematic^59,61^. We show for the first time that MSI cell lines harbouring *KMT2D*^LOF^ alterations are more susceptible to *WRN* KO than *KMT2D*^WT^ MSI cell lines, and *KMT2D*^LOF^ alterations co-occurred in WRN-dependent lines. Furthermore, we show that [AT]_n_ repeat elements were larger and more abundant than *KMT2D*^WT^ MSI cell lines. Given that WRN inhibitors are currently under preclinical development, our results thus provide the combination of MSI and *KMT2D* mutational status as potential biomarkers for future WRN-inhibitor treatments.

In *KMT2D*’s genetic network, we also identified regulators of the OXPHOS pathway, namely *NDUFB5^51^* (a priority class A SL candidate) and *OGT^115^* (a candidate co-essential gene and its encoded protein was detect in the ChIP-MS analysis), and showed that glycolytic genes were dysregulated in *KMT2D*^LOF^ TCGA-STAD cases. These results showing dysregulation of metabolic processes are consistent with the studies conducted in KMT2D-deficient mouse lung adenocarcinoma cell lines^53^ and in Kabuki patient-derived cells^54^. Interestingly, the OXPHOS pathway (NDUFB5 is a component of complex I), glycolysis, and OGT are known to contributes to the *de novo* pyrimidine synthesis^116,117^, a pathway that is garnering increasing attention as a therapeutic target^118^. A recent study by Gwynne et al.^117^ has shown that, in medulloblastoma, MYC (a downstream target of the MDM2/p53 pathway^77^) is stabilised by O-GlcNAcylation and that medulloblastoma cells were sensitive to inhibition of the *de novo* pyrimidine synthesis. Notably, our analyses also identified *MDM2* as a priority class A SL candidate and that *MDM2* and p53 downstream targets were dysregulated in *KMT2D*^LOF^ TCGA-COAD/READ cases. Taken together, we posit that *KMT2D*^LOF^ cases may also be more sensitive to *de novo* pyrimidine synthesis inhibitors than *KMT2D*^WT^ cases and that further studies are warranted to determine whether *KMT2D* mutations may be a biomarker for such targeted therapies.

Another important contribution of our study is our results further implicating KMT2D with a role in immune response regulation. MSI is a marker for ICI treatment; however, only ∼53% of MSI cases have an objective response to ICI treatment^119^, suggesting additional biomarkers are necessary to stratify MSI cases. Several biomarkers have been proposed for ICI treatment, including HRD^120^, *ARID1A*^LOF121^, and *SETD2*^LOF122^. Wang et al.^123^ also demonstrated that mice transplanted with KMT2D-deficient tumours were responsive to ICI treatment and that TMB was elevated in TCGA cases with *KMT2D*^LOF^. In this study, we show for the first time that candidate *KMT2D* SL interactors were involved in T-cell cytotoxicity and immune response regulation. Notably, we identified *TUBA1B* as a highly attractive SL candidate, where its overexpression has recently been implicated in favourable ICI treatment outcomes ^47^. We not only show that *TUBA1B* was overexpressed in *KMT2D*^LOF^ MSI cancers compared to *KMT2D*^WT^ MSI cancers in two cohorts, TCGA and POG, but we also show that several established ICI response markers are elevated in these cases. Previous studies have shown that a larger proportion of *KMT2D*^LOF^ cases had durable response to ICI treatment compared to *KMT2D*^WT^ cases^123,124^, and using TCGA and POG cohorts our results extend this notion to suggest that cases with MSI *KMT2D*^LOF^ cases would respond more favourably to ICI treatment than MSI *KMT2D*^WT^ cases. Altogether, these results strongly suggest that *KMT2D*^LOF^ MSI cancers may respond to ICI treatments, and we thus propose *KMT2D* as a possible biomarker to further stratify MSI cases for ICI treatment.

Our study of *in silico* genetic networks identified several potential roles for KMT2D and further implicated it in regulation of genomic stability and metabolism. We highlight cancer type-specific genetic interactors, namely *WRN*, *MDM2*, *NDUFB5*, and *TUBA1B*, which are promising targets of existing and in-development drugs. Furthermore, using cancer patient data, we provide evidence for *KMT2D*^LOF^ alterations as a potential biomarker for ICI treatment. Altogether, our work serves as an example for identifying novel functional associations and potential targeted treatment opportunities for cancers with tumour suppressor gene LOF alterations.

## Methods

### Generation of HEK KMT2D ^KO^ lines

The human embryonic kidney cell line HEK293A (provided by Dr. Gregg Morin, Genome Sciences Centre, BC Cancer and authenticated by Genetica DNA Laboratories, Cincinnati, OH) was cultured in Dulbecco’s modified Eagle’s Medium (DMEM; Life Technologies) supplemented with 10% (v/v) heat-inactivated fetal bovine serum (FBS; Life Technologies) in a 37°C incubator with 5% CO2 humidified atmosphere. Generation of HEK293A *KMT2D* knockout cells (herein referred to as *KMT2D*^KO^; *KMT2D*^KO1^ [D320] and *KMT2D*^KO3^ [D372]) were performed using CompoZr Knockout Zinc Finger Nucleases (CKOZFND14397, Sigma) targeting KMT2D exon 39 (Chr12: 49,427,414-49,427,455, NCBI build hg19). Each individual ZFN recognises an 18 bp DNA sequence on either side of a 6 bp spacer sequence where the heterodimerised FokI domains cleave the DNA. In total, the ZFN recognition and cleavage site (underlined) consists of 42 bases:

GGGATCCAGCCCCACCAG<u>AATGTT</u>GCTGTTGCTGCTGTTGGG These KMT2D-specific ZNF plasmids were co-transfected with a Hygro-RS-MLL2 ZNF reporter plasmid (ToolGen) into HEK293A. This reporter plasmid was used to assess ZFN efficiency^132^ through the co-expression of the mRFP gene, the ZNF recognition/cleavage sites and an out-of-frame Hygro-eGFP gene, which has a one in three chance of an in-frame repair following ZNF nuclease activity. Forty-eight hours post-transfection, the cells were single-cell sorted based on double RFP+/GFP+ expression using the FACSAriaII (BD Biosciences). Single cells were expanded and screened for clones bearing mutation of KMT2D in all alleles through immunoblotting, and frameshift mutations were confirmed through sequencing.

### ChIP-MS analysis

#### SILAC and ChIP

For heavy labelling, HEK-KMT2D^WT^ cells were grown in Dulbecco’s Modified Eagle Medium (DMEM) deficient in lysine and arginine (Thermo Fisher Scientific) supplemented with dialysed FBS (Thermo Fisher Scientific), MEM Non-Essential Amino Acids (Thermo Fisher Scientific), Lys-8 (CNLM-291-H-0.25, Cambridge Isotope Laboratories) Arg-10 (CNLM-539-H-0.25, Cambridge Isotope Laboratories) for 6 days before using the cells for ChIP. ChIP-MS was performed as described by Engelen et al. ^133^ with substitution of physical chromatin fragmentation (by sonication) with enzymatic treatment to preserve the integrity of high molecular weight proteins. Briefly, isolated nuclei were washed twice with 1X MNase Buffer (0.1% Na-Deoxycholate, 0.1%Triton X-100, CEF + NaButyrate) followed by centrifugation at 4°C for 5 minutes at 2800 rpm and supernatant discarded. Each pellet was resuspended in 3 pellet volumes of 1X MNase Buffer. Four pellet volumes of 2X MNase digestion buffer containing 1mM DTT and the MNase enzyme (diluted 1:100 in the total volume) were added. MNase digestion was performed at room temperature for 7 minutes and then stopped with EGTA to a final concentration of 20mM. MNase Buffer was added to a final concentration of 1X to help chromatin solubilisation on ice for 15 minutes. Digested chromatin was resuspended in IP buffer (200mM NaCl, 1mM EDTA, 0.15% NaDOC,1.5% Triton, 0.1% SDS) and samples were sonicated for 15 seconds (Covaris, Duty 10% - Intensity 4 - Burst 200 in 1 ml tubes) to break the nuclei. DNA fragments size was confirmed to be between 150 and 400 bp. Approximately 120 mg chromatin (as measured by Nanodrop at UV absorbance at 280 nm) were used for each sample and extracts from HEK-*KMT2D*^WT^ and HEK-*KMT2D*^KO1^ or HEK-*KMT2D*^KO3^ were mixed at one-to-one ratio. Two independent replicates for each *KMT2D* WT:KO mix underwent chromatin immunoprecipitation. Relative amounts of protein present in each sample were visualised on 10% precast SDS–PAGE gels (NuPage Invitrogen) and stained with colloidal Coomassie stain (Invitrogen) after cross linking reversal in the buffer containing 100mM NaHCO2, 2% SDS and followed by overnight incubation at 68℃. The following day the reaction was complemented with 20mM MgCl2 and 10ul of Benzonase nuclease (Millipore) to remove the DNA. Antibody used in the ChIP was against KMT2D (Sigma Prestige). For each ChIP, 50 μg antibody were crosslinked to 500 μl Protein A magnetic bead solution (15 mg beads, Life Technologies) with Dimethyl Pimelimidate (Sigma) to prevent the interference of immunoglobulin elution with the MS analysis. Crosslinked antibody–bead complexes were equilibrated in the buffer containing 20mM TrisHCl pH 8, 300mM NaCl, 2mM EDTA, 1mM EGTA, 0.2% NaDOC, 2% Triton X-100 and pre-cleared with 0.5 mg/ml of BSA, 0.2 mg/ml sonicated salmon sperm DNA for 1 h. The antibody–bead mixture was rotated overnight at 4°C. Beads were transferred to 1.5 ml no stick tubes (Alpha Laboratories) and washed following the protocol described by Engelen et al.^133^. The final three washes were performed with 0.2M HEPES pH 8.5 and samples eluted twice in 60 ul (each) of 50mM HEPES pH 8.5, 5mM dithiothreitol (DTT), 5% (weight per volume; w/v) SDS at 90 °C for 5 minutes on a thermocycler with shaking (800rpm). A small portion of the IP eluate (10%) was run on a polyacrylamide gel and stained with Silver staining to check the quality of the ChIP and reverse crosslinking. The remaining material was treated with 20mM chloroacetamide for 30 minutes at room temperature. Excess chloroacetamide was quenched through the addition of DTT to a final concentration of 40mM. Eluted proteins were then processed using a modified version of the SP3 protocol^134,135^. Specifically, 250µg of a combined and rinsed stock of Sera-Mag carboxylate-modified magnetic beads was combined with eluted proteins in a working volume of 100µL. Aggregation of proteins to the surface of the magnetic beads was driven via addition of acetonitrile to a final proportion of 80% (volume per volume; v/v). SP3 reactions were incubated for 5-minutes at room temperature, placed on a magnetic rack, and the supernatant was discarded. The beads were reconstituted in 800µL of 80% (v/v) ethanol, and placed on the magnetic rack again for supernatant removal. After supernatant removal, beads were reconstituted in 100mM HEPES pH 7.3 containing trypsin + rLysC (Promega) at a ratio of approximately 1:100µg (trypsin:sample) and incubated at 37°C for 18-hours in a ThermoMixer with shaking at 1,000rpm. Peptide-containing supernatants were recovered via centrifugation at 12,000g for 2-minutes along with use of a magnetic rack and then desalted using C18 SlitPlates (Glygen Scientific). Desalting was carried out by activating the column with methanol, rinsing with 0.1% (v/v) trifluoroacetic acid, sample binding, rinsing with 4% (v/v) methanol in 0.1% (v/v) formic acid, and final elution in 60% (v/v) methanol in 0.1% (v/v) formic acid. Desalted peptides were dried in a SpeedVac and reconstituted in a solution of 1% (v/v) dimethylsulfoxide (DMSO) and 1% (v/v) formic acid.

#### MS analysis

DDA-MS analysis was carried out on an Orbitrap Fusion instrument equipped with an EASY-nLC 1200 liquid chromatography (LC) system. For analysis, peptides were initially trapped on a pre-column (100µm x 3cm, 1.9µm C18 Reprosil-Pur Basic beads from Dr. Maisch) and then separated by an analytical column (100µm x 25cm, 1.9µm C18 Reprosil-Pur Basic beads) coupled to an in-house pulled tip electrospray emitter (∼20µm) with a 90-minute gradient at a flow rate of 400nL/min. Data acquisition on the Orbitrap Fusion used a standard data-dependent acquisition scheme. Specifically, the Orbitrap Fusion was globally set to use a positive ion spray voltage of 2200V, an ion transfer tube temperature of 275°C, a default charge state of 1, and an RF Lens setting of 60%. MS1 survey scans covered a mass range of 380-1500m/z at a resolution of 120,000 with an automatic gain control (AGC) target of 2e5 and a max injection time of 30ms. Precursors for tandem MS/MS (MS2) analysis were selected using monoisotopic precursor selection (‘Peptide’ mode), charge state filtering (2-4z), dynamic exclusion (20 ppm low and high, 30s duration), and an intensity threshold of 5e3. MS2 scans were carried out in the ion trap using the ‘Rapid’ scan mode, a fixed-first mass of 110m/z, a higher-energy collision dissociation setting of 32%, an AGC target of 1e4, a max injection time of 30ms, and an isolation window setting of 1m/z. The total allowable MS2 cycle time was set to 4s. MS1 data were acquired in profile mode, and MS2 in centroid. Resulting MS data were processed to obtain peptide and protein identifications using FragPipe (v17.1) with MSFragger (v3.4)^136^ and Philosopher (v4.1.1)^137^ with the default settings. Specifically, raw data were searched using MSFragger using the settings: 1. 20ppm precursor error; 2. 0.6 Daltons fragment error; 3. 2 missed cleavages; 4. +15.99@M, +42.01@N-term, +8.01@K, +10.00@R as variable modifications; 5. +57.02@C as a fixed modification; 6. Max precursor charge = 4. Data was searched against the Uniprot human database (version 2021-12) using a target-decoy strategy and filtered to provide an ∼1% FDR. Quantification of light and heavy peptide peaks was performed using the IonQuant node within FragPipe with the default settings. Proteins with a positive log_2_ fold change (mean HEK-KMT2D^WT^ replicates / mean HEK-KMT2D^LOF^ replicates) and at least one unique peptide in a replicate were considered candidate KMT2D interactors.

### Identifying KMT2D mutations across cancer types

Mutation annotation format (MAF) file from TCGA pan-cancer dataset, representing 10,217 tumour samples across 33 cancer types^35^, were downloaded from https://gdc.cancer.gov/about-data/publications/pancanatlas (mc3.v0.2.8.PUBLIC.maf; accessed on June 15th, 2022). Additionally, we downloaded mutation annotations from supplementary files from B-cell non-Hodgkin lymphoma (B-NHL) datasets (117 samples)^138–140^, medulloblastomas (MED) dataset (53 samples)^141^, and a small cell lung cancers (SCLC) dataset (110 samples)^142^, and annotated LOF mutations (nonsense mutations, frameshift insertions and deletions, and nonstop mutations). We used TCGA study abbreviations, except for B-NHL, which includes diffuse large B-cell lymphoma, follicular lymphoma, mantle cell lymphoma, and nodal marginal zone lymphomas, and COAD/READ, which includes colon and rectal adenocarcinomas. The following are the cancer type abbreviations used: adrenocortical carcinoma (ACC), bladder urothelial carcinoma (BLCA), B-cell non-Hodkin lymphoma (B-NHL), breast invasive carcinoma (BRCA), breast fibroepithelial (BRFE), cervical squamous cell carcinoma and endocervical adenocarcinoma (CESC), cholangiocarcinoma (CHOL), colon and rectal adenocarcinoma (COAD/READ), esophageal carcinoma (ESCA), glioblastoma multiforme (GBM), head and neck squamous cell carcinoma (HNSC), kidney chromophobe (KICH), kidney renal clear cell carcinoma (KIRC), kidney renal papillary cell carcinoma (KIRP), acute myeloid leukemia (LAML), brain lower grade glioma (LGG), liver hepatocellular carcinoma (LIHC), lung adenocarcinoma (LUAD), lung squamous cell carcinoma (LUSC), medulloblastoma (MED), mesothelioma (MESO), ovarian serous cystadenocarcinoma (OV), pancreatic adenocarcinoma (PAAD), pheochromocytoma and paraganglioma (PCPG), prostate adenocarcinoma (PRAD), sarcoma (SARC), small cell lung cancer (SCLC), skin cutaneous melanoma (SKCM), stomach adenocarcinoma (STAD), testicular germ cell tumors (TGCT), thyroid carcinoma (THCA), thymoma (THYM), uterine corpus endometrial carcinoma (UCEC), uterine carcinosarcoma (UCS), and uveal melanoma (UVM).

### In silico genetic network mapping

#### Essentiality network maps

Essentiality mapping and *in silico* genetic interaction screening were performed using our R software package GRETTA (0.99.2)^15^ with DepMap public release version 20Q1^143^. Briefly, co-essentiality mapping uses fitness effect scores derived from DepMap’s whole-genome CRISPR-Cas9 knockout screens, which targeted 18,333 genes in each of the 739 cancer cell lines. GRETTA calculated and ranked Pearson correlation coefficients between fitness scores of *KMT2D* and each screened gene. A gene pair was considered co-essential if its Pearson correlation p-value was < 0.05 and its coefficient was greater than the inflection point of the positive curve, and anti-essential if its p-value was < 0.05 and less than the inflection point of the negative curve.

#### Identifying DepMap cell lines

Using GRETTA (0.99.2)^15^ we queried *KMT2D* and *WRN* mutations from a total of 739 cancer cell lines in DepMap. These lines have WGS or whole-exome sequencing (WES) and RNA sequencing data, in addition to, genome-wide CRISPR-Cas9 KO screens^43,144^. Whole proteome quantification and global histone quantification were only available for a subset of 375 and 897 cancer cell lines, respectively^43,145^. The microsatellite status of DepMap cell lines was identified from supplementary files provided by Ghandi et al.^43^. As performed previously^6,15^, we used GRETTA to query the genomic data (MAF and copy number data files) to identify *KMT2D*^LOF^ and *WRN*^LOF^ mutant cancer cell lines (homozygous, trans-heterozygous, and heterozygous LOF mutants) and *KMT2D*^WT^ and *WRN*^WT^ control cell lines. Only trans-heterozygous lines (lines with more than one LOF mutation or a combination of LOF mutation and copy number loss) and heterozygous lines were available for the *KMT2D* pan-cancer group. Since trans-heterozygous LOF mutant groups are more likely to be KMT2D deficient than heterozygous lines^146^, we only used these lines in the *KMT2D*^LOF^ cell line group (16 lines). All LOF mutants (trans-heterozygous and heterozygous *KMT2D*^LOF^ lines) lines were considered in the *KMT2D* context-specific screens. For the pan-cancer *WRN* genetic screen, only heterozygous LOF mutants (176 lines) were available. To match DepMap cell lines to the appropriate cancer types, we used the “disease” and “disease subtype” columns provided in the cell line annotation file (sample_info.csv^43^) according to Supplementary Table S12. The normalised *KMT2D* TPM gene expression and protein expression values were extracted using GRETTA.

#### Differential lethality analysis

As described previously^6,15^, we used GRETTA to perform pairwise Mann-Whitney U tests comparing lethality probabilities for all 18,333 genes targeted in DepMap’s CRISPR-Cas9 KO screen between LOF mutant lines and WT lines to obtain p-values. P-values were adjusted for multiple testing using a permutation approach by randomly resampling 10,000 of the lethality probability scores. We used a threshold of adjusted p-value < 0.05 to identify candidate GIs of *KMT2D* in each screen, which were then prioritised (described in detail below). For visualisation, we calculated the genetic interaction score:

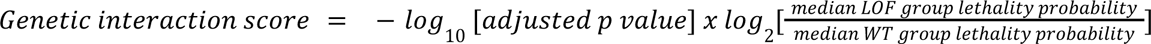

#### Prioritising candidate GIs

Candidate *KMT2D* interactors were prioritised according to their GI significance and their drug tractability, similar to the approaches used by Behan et al.^8^. Firstly, candidates were grouped into three tiers (tiers I, II, and III) based on their GI significance. Tier I candidates were the most significant group with adjusted Mann Whitney-U test p-values < 0.01, an absolute log_2_ fold change > 2 between median lethality probabilities (LOF group / WT group), and a minimum lethality probability of 0.5 in at least one group (WT or LOF). The following GI tiers II and III used progressively less stringent thresholds as shown in Figure 2D.

Next, candidates were grouped based on drug tractability (group I, II, and III). We used the Open Target Platform^45^, a database that contains a collection of tractability assessments based on sources, such as UniProt, HPA, PDBe, DrugEBIlity, ChEMBL, Pfam, InterPro, Complex Portal, DrugBank, Gene Ontology, and BioModels (https://platform-docs.opentargets.org/target/tractability; tractability_v23-02.tsv; accessed on February 15th, 2022), which is a strategy developed by Brown et al.^147^ and Schneider et al.^148^). Using this Open Target, we assessed whether the candidate GI encodes a protein for which there is already a known drug or, based on its protein structure, the likelihood of a drug being developed. Drug tractability group I contains GI candidates that are most tractable, and encodes proteins that are currently targetable using approved or phase I, II and II small molecule inhibitors or antibodies (buckets 1-3; documented on https://github.com/chembl/tractability_pipeline_v2/tree/master; accessed on accessed on February 15th, 2022). Group II contains targets that have high quality protein structure, ligand, and pocket annotations and cellular location annotation (buckets 4-6). Group III contains targets with proteins that have medium to low quality pocket annotation, cellular location annotation, or are known members of the druggable protein family (buckets 7-9).

Finally, GI candidates were prioritised by combining the GI significance and drug tractability assessments (class A, B, C, and D). With progressively less likelihood of targetable GIs, class A contained GIs with the highest significance with targetable proteins, and class D containing GI with the least significance with no evidence of tractability.

### Analysis of WRN dependency validated cell lines

Cancer cell lines that were validated to be WRN-(in)dependent by Behan et al.^8^, Chan et al.^59^, Kategaya et al.^58^, Lieb et al.^60^ and Picco et al.^62^, were identified and cross referenced to DepMap cell lines with WGS or WES annotation (32 cell lines). For each cell line, we annotated with reported WRN dependence and microsatellite status. Using DepMap WGS/WES annotations, we identified and annotated LOF alterations, including copy number loss, nonsense mutations, frameshift insertions and deletions, and nonstop mutations, affecting *KMT2D* and MMR genes (namely *EXO1*, *MBD4*, *MLH1*, *MSH2*, *MSH3*, *MSH5*, *MSH6*, *PMS1*, *PSM2*, and *RFC1*). Fisher’s exact tests were performed to determine co-occurrence between *KMT2D*/MMR gene LOF mutations and WRN dependence, and between *KMT2D* and MMR gene LOF alterations.

### Microsatellite repeat expansion analysis

I identified MSI DepMap cancer cell lines with WGS data that carried a WT *KMT2D* allele or LOF alterations (WT line: SKMEL2 and LOF lines: CCK81, LOVO, SW48, LS180, MFE319, and RKO) as well as two MSS lines (NCIH747 and COLO201; SRA project ID: PRJNA523380; SRA run IDs in Supplementary Table S5A). We downloaded the raw .fastq data using SRA-Toolkit (v2.11.3; https://trace.ncbi.nlm.nih.gov/Traces/sra/sra.cgi?view=software) and aligned to GRCh38 using bwa (v0.7.17)^149^. We used default ExpansionHunter Denovo (v0.9.0)^71^ settings to detect AT-motifs and filtered for regions that contained at least two in-repeat reads (i.e. one read pair maps within and the other outside of a repeat region).

### TCGA cohort analyses

#### Identifying *KMT2D* mutations

I used cBioPortal^150,151^ to download clinical (including MSIsensor, MANTIS, and TMB scores) and genomic data from TCGA Pan-Cancer Atlas Studies (33 cancer types). A TCGA case was considered to have MSI if the MSIsensor and MANTIS scores were >= 10 and >= 0.4, respectively, as previously determined^152,153^. We annotated cases with *KMT2D* LOF alterations as having nonsense mutation, frameshift insertions, frameshift deletions, or deep copy number loss. Cases with missense or silent mutations were excluded.

#### Mutational signature analysis

A matrix of 96 trinucleotide substitutions were downloaded from Alexandrov et al.^105^ at https://www.synapse.org/#!Synapse:syn11726618. The matrix was analysed in Python (v3.9) using the SigProfilerAssignment^154^ library to score single base pair substitution (SBS) signature activity scores with COSMIC (v.3.3), exome normalisation, and default settings.

#### Neoantigen and HRD scores and RNA-seq data

Neoantigen scores and HRD scores were downloaded from Thorsson et al.^96^ and RNA-seq count data were downloaded from UCSC Xena Hub^155^. Additional immune marker analyses are described below.

### POG cohort analyses

#### Ethics approval, consent to participate, and enrollment criteria

This work, involving POG program patients, was approved by the University of British Columbia–BC Cancer Research Ethics Board (H12-00137, H14-00681), and the POG program is registered under clinical trial number NCT02155621. All patients in this study gave informed written consent and were enrolled into POG as described previously, between July 2012 and May 2022^13,14,156^. The main objectives for our POG cohort analyses were to retrospectively validate TCGA cohort findings using whole-genome and whole-transcriptome analysis.

#### Tissue collection

Tissue collection and sequencing were performed as in Pleasance et al.^156^ for 820 patient samples. Tumour specimens were collected using needle core biopsies or tissue resection, snap frozen generally in optimal cutting temperature compound, reviewed by pathologists, and DNA and RNA libraries constructed and sequenced. Genome data were used to detect somatic alterations, including SNVs, copy number variants, loss of heterozygosity, and structural variants (such as gene fusions).

#### Whole-genome and transcriptome libraries

PCR-free genomic DNA libraries and poly-A selected RNA libraries were constructed and sequenced. Details can be found in Pleasance et al.^156^. In cases with RNA Integrity Numbers < 7.0, ribosomal RNA depletion RNA sequencing was employed as described in Pleasance et al. ^156^. Recent samples (since POG1046) were processed using an updated ribosomal RNA depletion RNA sequencing protocol, which is described below.

To remove cytoplasmic rRNA and mitochondrial ribosomal RNA (rRNA) species from total RNA, NEBNext rRNA Depletion Kit for Human/Mouse/Rat was used (NEB, E6310X). Enzymatic reactions were set-up in a 96-well plate (Thermo Fisher Scientific) on a Microlab NIMBUS liquid handler (Hamilton Robotics, USA). 150 ng of DNase treated total RNA in 6 µL was hybridized to rRNA probes in a 8.5 µL reaction. Heat-sealed plates were incubated at 95°C for 2 minutes followed by incremental reduction in temperature by 0.1°C per second to 22°C (730 cycles). The rRNA in DNA hybrids were digested using RNase H in a 11 µL reaction incubated in a thermocycler at 50°C for 30 minutes. To remove excess rRNA probes (DNA) and residual genomic DNA contamination, DNase I was added in a total reaction volume of 26 µL and incubated at 37°C for 30 minutes. RNA was purified using RNA MagClean DX beads (Aline Biosciences, USA) with 15 minutes of binding time, 7 minutes clearing on a magnet followed by two 70% ethanol washes, 5 minutes to air dry the RNA pellet and elution in 18µL DEPC water. The plate containing RNA was stored at -80°C prior to cDNA synthesis.

First-strand cDNA was synthesized from the purified RNA (minus rRNA) using the Maxima H Minus First Strand cDNA Synthesis kit (Thermo-Fisher, USA) and random hexamer primers at a concentration of 8 ng/µL along with a final concentration of 0.04 µg/µL Actinomycin D, followed by PCR Clean DX bead purification on a Microlab NIMBUS robot (Hamilton Robotics, USA). The second strand cDNA was synthesized following the NEBNext Ultra Directional Second Strand cDNA Synthesis protocol (NEB) that incorporates dUTP in the dNTP mix, allowing the second strand to be digested using USER^TM^ enzyme (NEB) in the post-adapter ligation PCR and thus achieving strand specificity.

cDNA was fragmented by Covaris LE220 sonication for 130 seconds (2 x 65seconds) at a “Duty cycle” of 30%, 450 Peak Incident Power (W) and 200 Cycles per Burst in a 96-well microTUBE Plate (P/N: 520078) to achieve 200-250 bp average fragment lengths. The paired-end sequencing library was prepared following Canada’s Micheal Smith Genome Sciences Centre strand-specific, plate-based library construction protocol on a Microlab NIMBUS robot (Hamilton Robotics, USA). Briefly, the sheared cDNA was subject to end-repair and phosphorylation in a single reaction using an enzyme premix (NEB) containing T4 DNA polymerase, Klenow DNA Polymerase and T4 polynucleotide kinase, incubated at 20°C for 30 minutes. Repaired cDNA was purified in 96-well format using PCR Clean DX beads (Aline Biosciences, USA), and 3’ A-tailed (adenylation) using Klenow fragment (3’ to 5’ exo minus) and incubation at 37°C for 30 minutes prior to enzyme heat inactivation. Ligation using TruSeq adapters at 20°C for 15 minutes. The adapter-ligated products were purified using PCR Clean DX beads followed by indexed PCR using NEBNext Ultra II Q5 Master Mix (NEB) with USER enzyme (1 U/µL, NEB) and dual indexed primer set. PCR parameters: 37°C for 15 minutes, 98°C for 1 minute followed by 13 cycles of 98°C 15 seconds, 65°C 30 seconds and 72°C 30 seconds, and then 72°C 5 minutes. The PCR products were purified and size-selected using a 1:1 PCR Clean DX beads-to-sample ratio (twice), and the eluted DNA quality was assessed with Caliper LabChip GX for DNA samples using the High Sensitivity Assay (PerkinElmer, Inc. USA) and quantified using a Quant-iT dsDNA High Sensitivity Assay Kit on a Qubit fluorometer (Invitrogen) prior to library pooling and size-corrected final molar concentration calculation for Illumina sequencing with paired-end 150 base reads.

#### Variant calling and *KMT2D* mutation selection

Somatic point mutations and small insertion and deletions are identified using Strelka 2.6.2^157^ and Mutect2 from GATK 4.0.10.0^158^. Variants from these tools are intersected using RTGTools^159^ to generate the calls as previously described^160^. We defined POG cases with *KMT2D* LOF alterations as those having somatic nonsense mutations, frameshift insertions, frameshift deletions, or deep copy number loss indicating likely homozygous deletions. Cases with missense or silent mutations were not considered and were excluded from the analysis entirely. POG cases with WT *KMT2D* alleles were defined as those with no somatic alteration and neutral copy number. Germline alterations were not considered in this study.

#### Immune marker analysis

Tumour mutation burden (TMB, mutations per megabase) is calculated by adding small mutations and small indels within the tumour sample generated in the above^160^ and dividing by the effective genome size (2934.876451 Mb). MSI were detected using genome data according to Pleasance et al.^156^ and HRDetect scores were calculated using a logistic regression model^161^. The model uses six mutation signatures associated with HRD as inputs into the algorithm: (i) Signature (SBS) 3, (ii) Signature (SBS) 8, (iii) SV signature 3, (iv) SV signature 5, (v) the HRD index, and (vi) the fraction of deletions with microhomology. All signatures were normalised and log transformed as previously described^156^. Additional immune marker analyses are described below.

### Survival analyses

Survival analyses were performed using R package ggsurvfit (v0.2.1). Differences in nonparametric survival functions were assessed across groups using log-rank tests with the ggsurvfit package.

### Additional immune marker analyses

Cytolytic activity scores were calculated as the geometric mean of *GZMA* and *PRF1* expression (FPKM), as performed by Rooney et al.^98^. M1/M2 macrophage scores were calculated according to Pender et al.^101^, as the mean expression (FPKM) of *CXCL11*, *IDO1*, *CCL19*, *CXCL9*, *PLA1A*, *LAMP3*, *CCR7*, *APOL6*, *CXCL10*, and *TNIP3*. IFN*γ* signature scores were calculated according to Ayers et al.^107^ by calculating the mean expression from ten genes associated with IFNγ signalling (*CCR5*, *CXCL10*, *CXCL11*, *CXCL9*, *GZMA*, *HLA-DRA, IDO1*, *IFNG, PRF1*, and *STAT1*). Immune cell infiltration proportion and absolute counts were calculated using the Cibersortx webportal (https://cibersortx.stanford.edu/) with default settings and the LM22 signature matrix file^109^. Immune checkpoint response scores were calculated using the PredictIO webportal (https://predictio.ca/), which uses a gene signature-based method to predict responders to ICI treatment^112^.

### Enrichment analysis and gene function annotation

We used clusterProfiler (v3.16.1)^162^ to identify pathway and protein complexes significantly enriched (q-value < 0.05) in the *KMT2D* essentiality network and protein interaction network, as well as to annotate functions of gene in the GI networks (unadjusted p-value < 0.05). Given many related terms with similar sets of genes associated with them, we grouped the terms based on their degree of similarity calculated with the Jaccard index, as performed in Takemon et al.^6^. We performed hierarchical clustering of the Jaccard index between pairs and the number of distinct clusters was determined using the gap statistic, which calculated the optimal number of clusters (up to 15 clusters) by iteratively bootstrapping 1000 times using the cluster (v2.1.4) package.

### Statistical analyses

All statistical analyses were conducted using R statistical software (v4.2.2)^163^ unless otherwise stated.

## Declarations

### Ethics approval and consent to participate

This study was approved by the UBC BC Cancer Research Ethics Board (H19-03010 and H20-02317). The work involving POG program patients was approved by the University of British Columbia–BC Cancer Research Ethics Board (H12-00137, H14-00681, H20-02317), and the POG program is registered under clinical trial number NCT02155621. All patients in this study gave informed written consent to participate and were enrolled into POG as described previously, between July 2012 and May 2022^13,14,156^.

### Consent for publication

Not applicable.

### Availability of data and materials

The scripts for reproducing the analysis, figures, and tables are available at GitHub (https://github.com/ytakemon/KMT2D_genetic_network_study). The GRETTA R package and DepMap data version 20Q1 used in this study are publicly available on the GRETTA GitHub repository (https://github.com/ytakemon/GRETTA). GRETTA has been archived with a citable DOI on Zenodo (https://doi.org/10.5281/zenodo.6940757) and a Singularity container has been made available on Sylabs (https://cloud.sylabs.io/library/ytakemon/gretta/gretta). ChIP-MS data have been deposited with the ProteomeXchange Consortium via the Proteomics Identification (PRIDE^164^) database (accession PXD048272). Genomic and transcriptomic sequence datasets, including metadata with library construction and sequencing approaches have been deposited at the European Genome–phenome Archive^165^ with accession numbers as listed in Supplemental Table S11. All other relevant data supporting the key findings of this study are available in Supplemental Tables.

### Author’s contributions

YT, AG, and MAM conceived the study and designed the experiments. YT, MAM, AG, CSH performed the experiments. EDP, AG, VC, KW, DLT, RDH, AJM, RAM, CH, KLM, EL, JN, HJL, DJR, SJJ, JL, and MAM provided resources. YT, EDP, VC, KW, AJM, RAM, CH, KLM, and EL curated the data. YT, CSH, and VG analysed the data. YT, AG, and MAM wrote the original draft of the manuscript. All authors edited and approved the manuscript. YT performed data visualisation. YT and JN provided project management. MAM provided supervision and acquired funding.

## Acknowledgements

This work would not be possible without the participation of patients and families, the POG team, the GSC platform, and the generous support of the BC Cancer Foundation, Genome British Columbia (project B20POG), The Terry Fox Research Institute, and the Terry Fox Foundation. MAM gratefully acknowledges the support of the Canada Research Chairs program, Genome BC, Genome Canada, and the University of British Columbia (UBC). YT is grateful for support from UBC’s 4-Year Fellowship, International Student Awards, and President’s Academic Excellence Initiative PhD Awards.

## Supplemental Figures

**Supplemental Figure S1.**
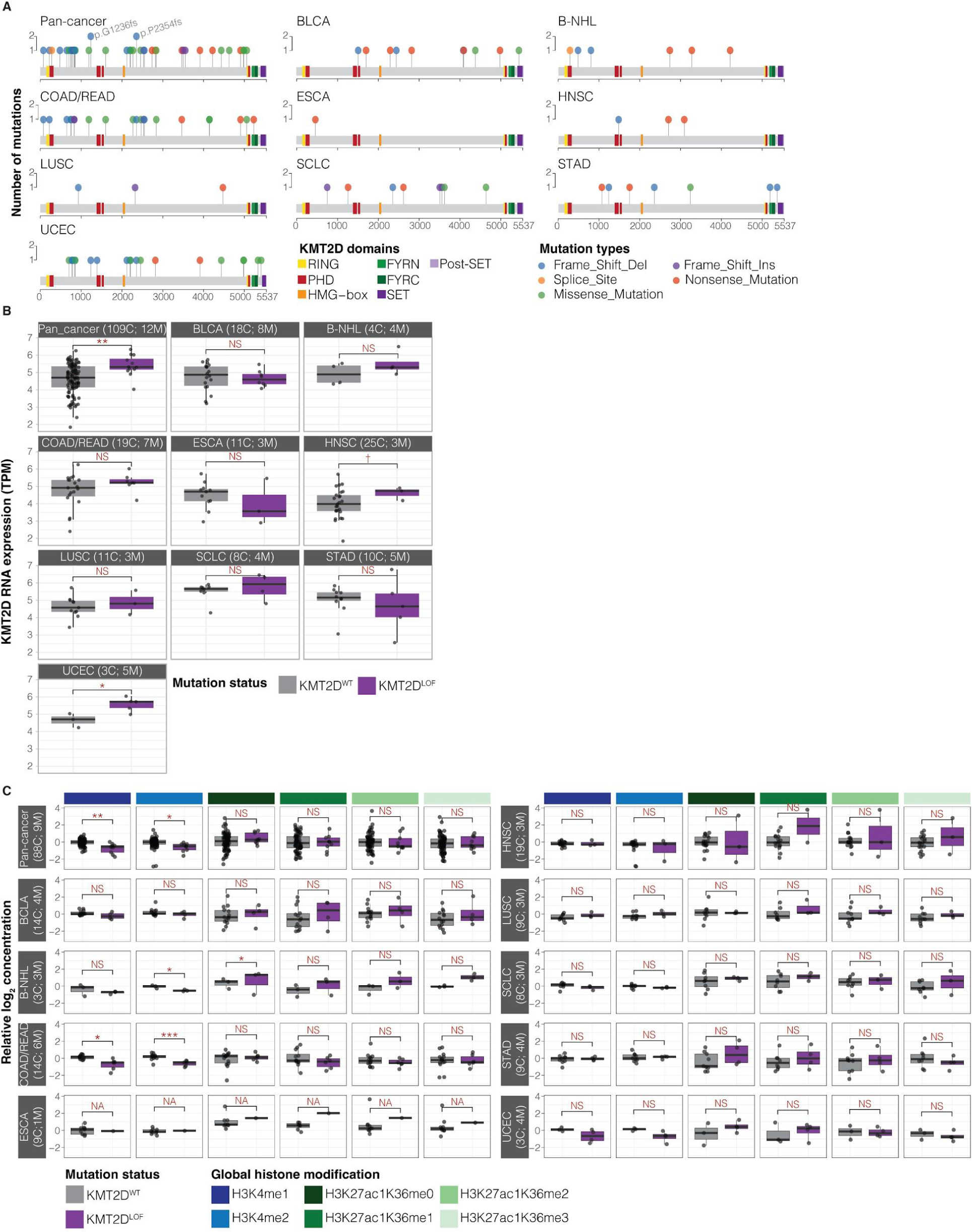
Characterisation of *KMT2D* mutations, expression, and global histone levels in DepMap cancer cell lines. **A.** Lollipop plots showing SNVs and small insertions and deletions in *KMT2D* identified in DepMap cell lines by cancer type. **B.** *KMT2D* mRNA expression in transcript per million (TPM) for *KMT2D*^WT^ and *KMT2D*^LOF^ DepMap cancer cell lines datasets. **C.** Relative concentration of global histone marks across cancer types. Benjamini Hochberg (BH)-corrected Welch’s t-test p-values ✝ < 0.1, * < 0.05, ** < 0.01, *** < 0.001 and NS > 0.1. NA indicates comparisons that are not analysed due to small sample size (N < 3).

**Supplemental Figure S2.**
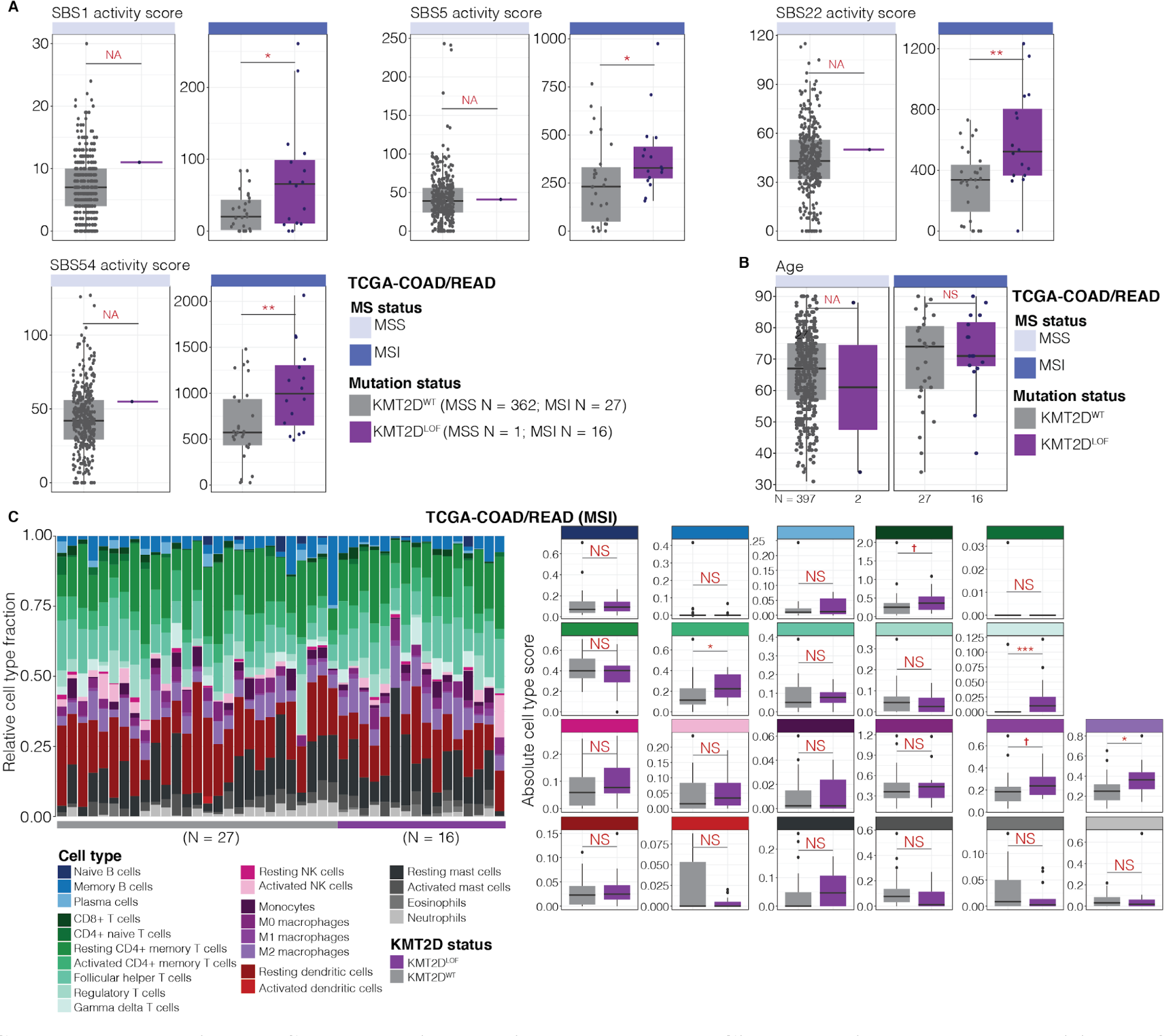
Mutational signatures and Cibersort immune composition of TCGA-COAD/READ cohorts. **A.** A comparison of single base pair substitution (SBS) mutational signatures. Uncorrected Welch’s t-test p-values * < 0.05 and ** < 0.01, and not analysed (NA) when sample size was < 3. Not significant after multiple-testing correction (BH-corrected p-value > 0.05). **B.** A comparison of chronological age. Welch’s t-test p-values NS > 0.1 and not analysed (NA) when sample size was < 3. **C.** Distribution of relative cell type fractions (left) and a comparison of absolute cell type fractions (right) calculated by Cibersortx in TCGA MSI COAD/READ cohort. Relative cell type fractions are shown in the left panels. BH-corrected Welch’s t-test p-values … < 0.1, * < 0.05, ** < 0.01, *** < 0.001, and NS > 0.1.

**Supplemental Figure S3.**
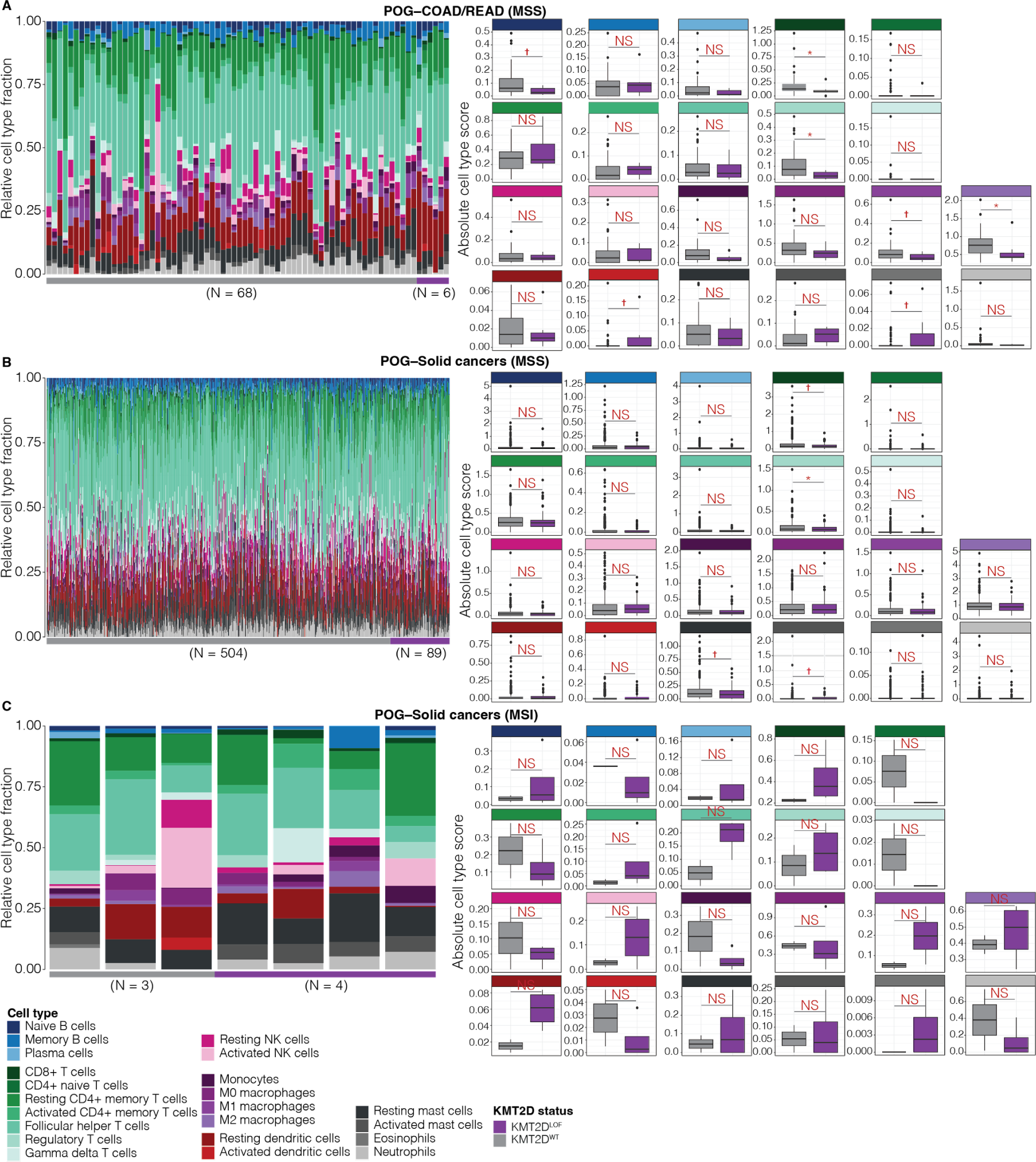
Cibersort immune composition of POG cohorts. **A-C.** Distribution of relative cell type fractions (left) and a comparison of absolute cell type fractions (right) calculated by Cibersortx in POG MSI-COAD/READ (A), MSS-solid cancer (B), and MSI-solid cancer (C) cohorts. Relative cell type fractions are shown in the left panels. BH-corrected Welch’s t-test p-values ✝ < 0.1, * < 0.05, and NS > 0.05.

## Supplemental Tables

**Supplemental Table S1: *KMT2D* essentiality network results.**

**A.** Ranked list of genes in the *KMT2D* essentiality network. GeneName: hugo gene symbol. GeneNameID: hugo gene symbol separated by NCBI gene id. Estimate: Pearson’s correlation coefficient. P.value: p-value derived from Pearson’s correlation test. Rank: rank in list determined by order of estimate. Candidate_inflection: TRUE if a gene’s Pearson’s correlation coefficient is past the inflection point of the positive/negative curve (see Methods). Chr: chromosome location of gene. Gene_start_bp: a gene’s start position. Gene_end_bp: a gene’s end position. Karyotype_band: the karyotype band in which the gene is found. **B.** Enriched GO biological processes found among co-/anti-essential genes. Results are outputs from ClusterProfiler. **C.** Clustered GO terms of co-essential genes.

**Supplemental Table S2: KMT2D ChIP-MS results.**

**A.** KMT2D ChIP-MS detected proteins. Accession: uniprot id. Gene: hugo gene symbol. LogFC_HL_kmt2d_R1KO1: log_10_ fold change (heavy isotope/light isotope) of replicate 1 of HEK-*KMT2D*^KO1^. LogFC_HL_kmt2d_R2KO1: log_10_ fold change (heavy isotope/light isotope) of replicate 2 of HEK-*KMT2D*^KO1^. LogFC_HL_kmt2d_R1KO3: log_10_ fold change (heavy isotope/light isotope) of replicate 1 of HEK-*KMT2D*^KO3^. LogFC_HL_kmt2d_R2KO3: log_10_ fold change (heavy isotope/light isotope) of replicate 2 of HEK-*KMT2D*^KO3^. Num_reps_with_peptides: number of replicates with peptides detected. **B.** Enriched GO biological processes found among detected proteins. Results are outputs from ClusterProfiler. **C.** Clustered GO terms. **D.** A list of KMT2D interactors that overlap with *KMT2D* co-/anti-essential genes.

**Supplemental Table S3: Results GI predictions using GRETTA.**

**A**. List of DepMap cancer cell lines by KMT2D^WT^/KMT2D^LOF^ genotypes and cancer type. DepMap_ID: DepMap cancer cell lines IDs. stripped_cell_line_name: cell line names provided by DepMap without special characters or spaces. disease: cancer type assigned by DepMap. disease_subtype: cancer subtype assigned by Depmap. TCGA_type: matched TCGA cancer type abbreviation (see Methods for details). Total_mutations: total number of mutations detected (LOF and non-LOF). LOF_mutations: number of LOF SNVs and insertion deletions detected (see Methods). CN_status: copy number status. Group: *KMT2D* group assigned (Control refers to the *KMT2D*^WT^ group). **B.** All *KMT2D* GI prediction results by cancer context. Cancer type context: cancer context *KMT2D* GI screens were performed in. GeneNameID: hugo gene name with Entrez gene ID. GeneNames: gene name without Entrez gene ID. Control_median: median lethality probability of *KMT2D*^WT^ cell line group. Mutant_median: median of *KMT2D*^LOF^ group lethality probabilities. Control_sd: standard deviation of *KMT2D*^WT^ lethality probabilities. Mutant_sd: standard deviation of *KMT2D*^LOF^ lethality probabilities. Control_iqr: interquartile range of *KMT2D*^WT^ lethality probabilities. Mutant_iqr: interquartile range of *KMT2D*^LOF^ lethality probabilities. Pval: uncorrected Mann Whitney U-test p-values comparing *KMT2D*^WT^ and *KMT2D*^WT^ lethality probabilities. log2FC_by_median: log2-transformed fold change of *KMT2D*^LOF^ over *KMT2D*^WT^ lethality probabilities. CliffDelta: cliff’s delta effect size comparing *KMT2D*^LOF^ and *KMT2D*^WT^ lethality probabilities. Dip_pval: hartigan’s dip test p-value to test unimodality of lethality probabilities. Interaction_score: GI interaction score for visualisation (see Methods). Adj_pval: Mann Whitney U-test p-values adjusted for multiple testing using permutation test with 10,000 random samplings. GI_direction: whether a GI is alleviating (AL) or synthetic lethal (SL). GI_tiers: GI prediction tier assigned to GI. **C.** Prioritisation classification assigned to significant GIs (GI prediction tiers I, II, and III). First 17 column names are the same as A. Drug group: drug tractability group assigned to GI. Priority_class: priority class assigned to GI.

**Supplemental Table S4: List of DepMap cancer cell lines in *WRN*^WT^/*WRN*^LOF^ groups.**

**A.** DepMap_ID: DepMap cancer cell lines IDs. stripped_cell_line_name: cell line names provided by DepMap without special characters or spaces. disease: cancer type assigned by DepMap. disease_subtype: cancer subtype assigned by Depmap. MS_status: inferred microsatellite status by Ghandi et al. (2019). Group: *WRN* group assigned (Control refers to the *WT*^WT^ group; pan-cancer). Total_mutations: total number of mutations detected (LOF and non-LOF). LOF_mutations: number of LOF SNVs and insertion deletions detected (see Methods). CN_status: copy number status. KMT2D_prob: lethality probability when *KMT2D* is knocked out in the cell line. **B.** DepMap cancer cell lines that were previously validated to be dependent or independent of WRN. DepMap_ID: DepMap cancer cell lines IDs. stripped_cell_line_name: Cell line names provided by DepMap without special characters or spaces. disease: cancer type assigned by DepMap. Author: first author of study. WRN_dependence: whether authors showed WRN dependence. TRUE indicates KO of WRN was lethal to cell lines and FALSE indicates that cells were viable when WRN was knocked out. **C.** KMT2D and MMR mutations detected by DepMap WGS/WES characterization in cell lines from B.

**Supplemental Table S5: *WRN* KO validated cell lines**

**A.** List of DepMap cell lines previously for WRN survival dependence using *WRN* KO assays. DepMap_ID: DepMap cell line id. Cell_line_name: Name of cell line. Author: name of publication *WRN* KO assay was performed in. WRN_dependence: outcome of *WRN* KO (TRUE indicates that WRN was essential for survival and FALSE indicates that it was not). **B.** Mutations found in MMR genes and *KMT2D* in cell lines in A.

**Supplemental Table S6: Characterising [AT]_n_ motif expansion in cancer cell lines.**

**A.** All MSI DepMap cancer cell lines with WGS data and three MSS DepMap cell lines with WGS data for comparison. Cell_line_stripped_name: cell line names provided by DepMap without special characters or spaces. disease: cancer type assigned by DepMap. MSI_status: inferred microsatellite status by Ghandi et al. (2019). KMT2D_group: *KMT2D* group assigned to a cell line. SRA_project_ID: NCBI SRA project ID for raw WGS data. SRA_Run_ID: NCBI SRA run ID for raw WGS data. Instrument: sequencer used to perform WGS. **B.** Profiles of [AT]_n_ motifs generated using ExpansionHunter Denovo. Cell_line_name: cell line names provided by DepMap without special characters or spaces. disease: cancer type assigned by DepMap. SRA_id: NCBI SRAN run ID. group: MSI status and KMT2D group of cell line. Contig: chromosome location of motif. Start: start site (bp) of motif. End: end site (bp) of motif. Motif: motif type. Num_anc_irrs: number of anchored in-repeat reads (i.e. one read pair maps within and the other outside of a repeat region; see Methods). norm_num_anc_irrs: normalised number of anchored in-repeat reads. Het_str_size: estimated motif repeat size.

**Supplemental Table S7: Expression of *MDM2*, *TP53*, and TP53-downstream targets in TCGA-COAD/READ cases.**

**A.** TCGA-COAD/READ cohort KMT2D and TP53 mutations and expression of *MDM2*, *TP53*, and TP53-downstream targets. mut: WT, LOF, or Other mutation called. cn: copy number status, tpm: transcript per million.

**Supplemental Table S8: Expression of *NDUFB5, PER2,* and PER2-downstream targets in TCGA-STAD cases.**

**A.** TCGA-STAD cohort *NDUFB5*, *PER2*, and PER2-target gene expression. tpm: transcript per million (see Methods section).

**Supplemental Table S9: Profiles of *KMT2D* mutations and immune markers in TCGA-COAD/READ cases.**

**A.** TCGA-COAD/READ clinical data and immune marker profiles. OS: overall survival. Fpkm: fragments per kilobase of transcript per million mapped reads. B_cells_naive to absolute_score_sig_score columns are immune cell composition score outputs from Cibersort. PredictIO_score: immune checkpoint response score (see Methods).

**Supplemental Table S10: Age and mutational signature activity scores of TCGA-COAD/READ cases.**

**Supplemental Table S11: Profiles of *KMT2D* mutations and immune markers in POG cases.**

**A.** POG solid cancer clinical data and immune marker profiles. OS: overall survival. censor_status: 1-living, 2-deceased. Fpkm: fragments per kilobase of transcript per million mapped reads. B_cells_naive to absolute_score_sig_score columns are immune cell composition score outputs from Cibersort. PredictIO_score: immune checkpoint response score (see Methods section).

**Supplemental Table S12: TCGA cancer type assigned to DepMap disease types.**

## References

1. Morris, L. G. T. & Chan, T. A. Therapeutic targeting of tumor suppressor genes. Cancer 121, 1357–1368 (2015).

2. Wang, T. et al. Gene Essentiality Profiling Reveals Gene Networks and Synthetic Lethal Interactions with Oncogenic Ras. Cell 168, 890–903 e15 (2017).

3. Pan, J. et al. Interrogation of Mammalian Protein Complex Structure, Function, and Membership Using Genome-Scale Fitness Screens. Cell Syst 6, 555–568.e7 (2018).

4. Kim, E. et al. A network of human functional gene interactions from knockout fitness screens in cancer cells. Life Sci Alliance 2, (2019).

5. Wainberg, M. et al. A genome-wide atlas of co-essential modules assigns function to uncharacterized genes. Nat. Genet. 53, 638–649 (2021).

6. Takemon, Y. et al. Multi-Omic Analysis of CIC’s Functional Networks Reveals Novel Interaction Partners and a Potential Role in Mitotic Fidelity. Cancers 15, 2805 (2023).

7. Mani, R., St Onge, R. P., Hartman, J. L., 4th, Giaever, G. & Roth, F. P. Defining genetic interaction. Proc. Natl. Acad. Sci. U. S. A. 105, 3461–3466 (2008).

8. Behan, F. M. et al. Prioritization of cancer therapeutic targets using CRISPR–Cas9 screens. Nature 568, 511–516 (2019).

9. De Kegel, B., Quinn, N., Thompson, N. A., Adams, D. J. & Ryan, C. J. Comprehensive prediction of synthetic lethality between paralog pairs in cancer cell lines. Cold Spring Harbor Laboratory 2020.12.16.423022 (2020) doi:10.1101/2020.12.16.423022.

10. Jiang, M., Instrell, R., Saunders, B., Berven, H. & Howell, M. Tales from an academic RNAi screening facility; FAQs. Brief. Funct. Genomics 10, 227–237 (2011).

11. Bock, C. et al. High-content CRISPR screening. Nature Reviews Methods Primers 2, 1–23 (2022).

12. Kandoth, C. et al. Mutational landscape and significance across 12 major cancer types. Nature 502, 333–339 (2013).

13. Laskin, J., et al. Lessons learned from the application of whole-genome analysis to the treatment of patients with advanced cancers. Cold Spring Harb Mol Case Stud 1, a000570 (2015).

14. Pleasance, E. et al. Pan-cancer analysis of advanced patient tumors reveals interactions between therapy and genomic landscapes. Nature Cancer 1, 452–468 (2020).

15. Takemon, Y. & Marra, M. A. GRETTA: an R package for mapping in silico genetic interaction and essentiality networks. Bioinformatics 39, (2023).

16. Fagan, R. J. & Dingwall, A. K. COMPASS Ascending: Emerging clues regarding the roles of MLL3/KMT2C and MLL2/KMT2D proteins in cancer. Cancer Lett. 458, 56–65 (2019).

17. Cho, Y.-W., Hong, S. & Ge, K. Affinity purification of MLL3/MLL4 histone H3K4 methyltransferase complex. Methods Mol. Biol. 809, 465–472 (2012).

18. Cho, Y.-W. et al. PTIP associates with MLL3- and MLL4-containing histone H3 lysine 4 methyltransferase complex. J. Biol. Chem. 282, 20395–20406 (2007).

19. Hong, S. et al. Identification of JmjC domain-containing UTX and JMJD3 as histone H3 lysine 27 demethylases. Proc. Natl. Acad. Sci. U. S. A. 104, 18439–18444 (2007).

20. Kantidakis, T. et al. Mutation of cancer driver MLL2 results in transcription stress and genome instability. Genes Dev. 30, 408–420 (2016).

21. Shang, J.-Y. et al. COMPASS functions as a module of the INO80 chromatin remodeling complex to mediate histone H3K4 methylation in Arabidopsis. Plant Cell 33, 3250–3271 (2021).

22. Mayayo-Peralta, I. et al. PAXIP1 and STAG2 converge to maintain 3D genome architecture and facilitate promoter/enhancer contacts to enable stress hormone-dependent transcription. Nucleic Acids Res. (2023) doi:10.1093/nar/gkad267.

23. Liu, B. & Li, Z. PTIP-Associated Protein 1: More Than a Component of the MLL3/4 Complex. Front. Genet. 13, 889109 (2022).

24. Hamadeh, Z. & Lansdorp, P. RECQL5 at the Intersection of Replication and Transcription. Front Cell Dev Biol 8, 324 (2020).

25. Morrison, A. J. & Shen, X. Chromatin remodelling beyond transcription: the INO80 and SWR1 complexes. Nat. Rev. Mol. Cell Biol. 10, 373–384 (2009).

26. Tothova, Z., et al. Cohesin mutations alter DNA damage repair and chromatin structure and create therapeutic vulnerabilities in MDS/AML. JCI Insight 6, (2021).

27. Froimchuk, E., Jang, Y. & Ge, K. Histone H3 lysine 4 methyltransferase KMT2D. Gene 627, 337–342 (2017).

28. Konzman, D. et al. O-GlcNAc: Regulator of Signaling and Epigenetics Linked to X-linked Intellectual Disability. Front. Genet. 11, 605263 (2020).

29. Liu, C. & Li, J. O-GlcNAc: A Sweetheart of the Cell Cycle and DNA Damage Response. Front. Endocrinol. 9, 415 (2018).

30. Gondane, A. et al. O-GlcNAc transferase couples MRE11 to transcriptionally active chromatin to suppress DNA damage. J. Biomed. Sci. 29, 13 (2022).

31. Cortez, D. Replication-Coupled DNA Repair. Mol. Cell 74, 866–876 (2019).

32. Pećina-Šlaus, N., Kafka, A., Salamon, I. & Bukovac, A. Mismatch Repair Pathway, Genome Stability and Cancer. Front Mol Biosci 7, 122 (2020).

33. Mondal, G., Stevers, M., Goode, B., Ashworth, A. & Solomon, D. A. A requirement for STAG2 in replication fork progression creates a targetable synthetic lethality in cohesin-mutant cancers. Nat. Commun. 10, 1686 (2019).

34. Saponaro, M. et al. RECQL5 controls transcript elongation and suppresses genome instability associated with transcription stress. Cell 157, 1037–1049 (2014).

35. Ellrott, K. et al. Scalable Open Science Approach for Mutation Calling of Tumor Exomes Using Multiple Genomic Pipelines. Cell Syst 6, 271–281.e7 (2018).

36. Rossi, A. et al. Genetic compensation induced by deleterious mutations but not gene knockdowns. Nature 524, 230–233 (2015).

37. Rouf, M. A. et al. The recent advances and future perspectives of genetic compensation studies in the zebrafish model. Genes Dis 10, 468–479 (2023).

38. Hu, D. et al. The MLL3/MLL4 branches of the COMPASS family function as major histone H3K4 monomethylases at enhancers. Mol. Cell. Biol. 33, 4745–4754 (2013).

39. Lee, J.-E. et al. H3K4 mono- and di-methyltransferase MLL4 is required for enhancer activation during cell differentiation. Elife 2, e01503 (2013).

40. Dhar, S. S. et al. MLL4 Is Required to Maintain Broad H3K4me3 Peaks and Super-Enhancers at Tumor Suppressor Genes. Mol. Cell 70, 825–841.e6 (2018).

41. Ortega-Molina, A. et al. The histone lysine methyltransferase KMT2D sustains a gene expression program that represses B cell lymphoma development. Nat. Med. 21, 1199–1208 (2015).

42. Zhang, J. et al. Disruption of KMT2D perturbs germinal center B cell development and promotes lymphomagenesis. Nat. Med. 21, 1190–1198 (2015).

43. Ghandi, M. et al. Next-generation characterization of the Cancer Cell Line Encyclopedia. Nature 569, 503–508 (2019).

44. Henkel, L., Rauscher, B. & Boutros, M. Context-dependent genetic interactions in cancer. Curr. Opin. Genet. Dev. 54, 73–82 (2019).

45. Ochoa, D. et al. The next-generation Open Targets Platform: reimagined, redesigned, rebuilt. Nucleic Acids Res. 51, D1353–D1359 (2023).

46. Hou, H., Sun, D. & Zhang, X. The role of MDM2 amplification and overexpression in therapeutic resistance of malignant tumors. Cancer Cell Int. 19, 216 (2019).

47. Hu, X. et al. Tubulin Alpha 1b Is Associated with the Immune Cell Infiltration and the Response of HCC Patients to Immunotherapy. Diagnostics (Basel) 12, (2022).

48. Hu, J. et al. Dynamic Network Biomarker of Pre-Exhausted CD8+ T Cells Contributed to T Cell Exhaustion in Colorectal Cancer. Front. Immunol. 12, 691142 (2021).

49. Eischen, C. M. Role of Mdm2 and Mdmx in DNA repair. J. Mol. Cell Biol. 9, 69–73 (2017).

50. Kim, J. M. Molecular Link between DNA Damage Response and Microtubule Dynamics. Int. J. Mol. Sci. 23, (2022).

51. Stroud, D. A. et al. Accessory subunits are integral for assembly and function of human mitochondrial complex I. Nature 538, 123–126 (2016).

52. Padavannil, A., Ayala-Hernandez, M. G., Castellanos-Silva, E. A. & Letts, J. A. The Mysterious Multitude: Structural Perspective on the Accessory Subunits of Respiratory Complex I. Front Mol Biosci 8, 798353 (2021).

53. Alam, H. et al. KMT2D Deficiency Impairs Super-Enhancers to Confer a Glycolytic Vulnerability in Lung Cancer. Cancer Cell 37, 599–617.e7 (2020).

54. Pacelli, C. et al. Loss of Function of the Gene Encoding the Histone Methyltransferase KMT2D Leads to Deregulation of Mitochondrial Respiration. Cells 9, (2020).

55. Futami, K. & Furuichi, Y. RECQL1 and WRN DNA repair helicases: potential therapeutic targets and proliferative markers against cancers. Front. Genet. 5, 441 (2014).

56. Mukherjee, S. et al. Werner Syndrome Protein and DNA Replication. Int. J. Mol. Sci. 19, (2018).

57. Marabitti, V. et al. R-Loop-Associated Genomic Instability and Implication of WRN and WRNIP1. Int. J. Mol. Sci. 23, (2022).

58. Kategaya, L., Perumal, S. K., Hager, J. H. & Belmont, L. D. Werner Syndrome Helicase Is Required for the Survival of Cancer Cells with Microsatellite Instability. iScience 13, 488–497 (2019).

59. Chan, E. M. et al. WRN helicase is a synthetic lethal target in microsatellite unstable cancers. Nature 568, 551–556 (2019).

60. Lieb, S. et al. Werner syndrome helicase is a selective vulnerability of microsatellite instability-high tumor cells. Elife 8, (2019).

61. van Wietmarschen, N. et al. Repeat expansions confer WRN dependence in microsatellite-unstable cancers. Nature 586, 292–298 (2020).

62. Picco, G. et al. Werner Helicase Is a Synthetic-Lethal Vulnerability in Mismatch Repair-Deficient Colorectal Cancer Refractory to Targeted Therapies, Chemotherapy, and Immunotherapy. Cancer Discov. (2021) doi:10.1158/2159-8290.CD-20-1508.

63. Mullard, A. What’s next for the synthetic lethality drug discovery engine? Nat. Rev. Drug Discov. 21, 477–479 (2022).

64. Morales-Juarez, D. A. & Jackson, S. P. Clinical prospects of WRN inhibition as a treatment for MSI tumours. npj Precision Oncology 6, 1–6 (2022).

65. Kim, T. Y. et al. Substrate trapping proteomics reveals targets of the βTrCP2/FBXW11 ubiquitin ligase. Mol. Cell. Biol. 35, 167–181 (2015).

66. Huttlin, E. L. et al. Architecture of the human interactome defines protein communities and disease networks. Nature 545, 505–509 (2017).

67. Chammas, P., Mocavini, I. & Di Croce, L. Engaging chromatin: PRC2 structure meets function. Br. J. Cancer 122, 315–328 (2020).

68. Liu, X. et al. Structural insights into dimethylation of 12S rRNA by TFB1M: indispensable role in translation of mitochondrial genes and mitochondrial function. Nucleic Acids Res. 47, 7648–7665 (2019).

69. Mu, J. et al. Mitochondrial transcription factor B1 promotes the progression of hepatocellular carcinoma via enhancing aerobic glycolysis. J. Cell Commun. Signal. 16, 223–238 (2022).

70. Velho, S., Fernandes, M. S., Leite, M., Figueiredo, C. & Seruca, R. Causes and consequences of microsatellite instability in gastric carcinogenesis. World J. Gastroenterol. 20, 16433–16442 (2014).

71. Dolzhenko, E. et al. ExpansionHunter Denovo: a computational method for locating known and novel repeat expansions in short-read sequencing data. Genome Biol. 21, 102 (2020).

72. Konopleva, M. et al. MDM2 inhibition: an important step forward in cancer therapy. Leukemia 34, 2858–2874 (2020).

73. Sugano, N. et al. MDM2 gene amplification in colorectal cancer is associated with disease progression at the primary site, but inversely correlated with distant metastasis. Genes Chromosomes Cancer 49, 620–629 (2010).

74. Frum, R. A. et al. The human oncoprotein MDM2 induces replication stress eliciting early intra-S-phase checkpoint response and inhibition of DNA replication origin firing. Nucleic Acids Res. 42, 926–940 (2014).

75. Lee, J. et al. A tumor suppressive coactivator complex of p53 containing ASC-2 and histone H3-lysine-4 methyltransferase MLL3 or its paralogue MLL4. Proc. Natl. Acad. Sci. U. S. A. 106, 8513–8518 (2009).

76. Rahnamoun, H. et al. Mutant p53 regulates enhancer-associated H3K4 monomethylation through interactions with the methyltransferase MLL4. J. Biol. Chem. 293, 13234–13246 (2018).

77. Rozan, L. M. & El-Deiry, W. S. p53 downstream target genes and tumor suppression: a classical view in evolution. Cell Death Differ. 14, 3–9 (2007).

78. Sullivan, K. D., Galbraith, M. D., Andrysik, Z. & Espinosa, J. M. Mechanisms of transcriptional regulation by p53. Cell Death Differ. 25, 133–143 (2018).

79. Farhang Ghahremani, M., et al. p53 promotes VEGF expression and angiogenesis in the absence of an intact p21-Rb pathway. Cell Death Differ. 20, 888–897 (2013).

80. Haupt, S., Gamell, C., Wolyniec, K. & Haupt, Y. Interplay between p53 and VEGF: how to prevent the guardian from becoming a villain. Cell death and differentiation vol. 20 852–854 (2013).

81. Maitituoheti, M. et al. Enhancer Reprogramming Confers Dependence on Glycolysis and IGF Signaling in KMT2D Mutant Melanoma. Cell Rep. 33, 108293 (2020).

82. Zhao, X., Tian, Z. & Liu, L. circATP2B1 Promotes Aerobic Glycolysis in Gastric Cancer Cells Through Regulation of the miR-326 Gene Cluster. Front. Oncol. 11, 628624 (2021).

83. Dogan, S. A. et al. Tissue-specific loss of DARS2 activates stress responses independently of respiratory chain deficiency in the heart. Cell Metab. 19, 458–469 (2014).

84. Kim, H.-J. & Barrientos, A. MTG1 couples mitoribosome large subunit assembly with intersubunit bridge formation. Nucleic Acids Res. 46, 8435–8453 (2018).

85. Fernandez-Vizarra, E. & Zeviani, M. Mitochondrial disorders of the OXPHOS system. FEBS Lett. 595, 1062–1106 (2021).

86. Sharoyko, V. V. et al. Loss of TFB1M results in mitochondrial dysfunction that leads to impaired insulin secretion and diabetes. Hum. Mol. Genet. 23, 5733–5749 (2014).

87. Center for Drug Evaluation & Research. FDA grants accelerated approval to pembrolizumab for first tissue/site agnostic indication. U.S. Food and Drug Administration https://www.fda.gov/drugs/resources-information-approved-drugs/fda-grants-accelerated-approval-pembrolizumab-first-tissuesite-agnostic-indication (2019).

88. Marcus, L., Lemery, S. J., Keegan, P. & Pazdur, R. FDA Approval Summary: Pembrolizumab for the Treatment of Microsatellite Instability-High Solid Tumors. Clin. Cancer Res. 25, 3753–3758 (2019).

89. Diaz, L. A., Jr et al. Pembrolizumab versus chemotherapy for microsatellite instability-high or mismatch repair-deficient metastatic colorectal cancer (KEYNOTE-177): final analysis of a randomised, open-label, phase 3 study. Lancet Oncol. 23, 659–670 (2022).

90. Ardeshir-Larijani, F. et al. KMT2D Mutation Is Associated With Poor Prognosis in Non-Small-Cell Lung Cancer. Clin. Lung Cancer 19, e489–e501 (2018).

91. Ferrero, S. et al. KMT2D mutations and TP53 disruptions are poor prognostic biomarkers in mantle cell lymphoma receiving high-dose therapy: a FIL study. Haematologica 105, 1604–1612 (2020).

92. Li, Q. et al. Plasma circulating tumor DNA assessment reveals KMT2D as a potential poor prognostic factor in extranodal NK/T-cell lymphoma. Biomark Res 8, 27 (2020).

93. Morcillo-Garcia, S. et al. Genetic mutational status of genes regulating epigenetics: Role of the histone methyltransferase KMT2D in triple negative breast tumors. PLoS One 14, e0209134 (2019).

94. Subbiah, V., Solit, D. B., Chan, T. A. & Kurzrock, R. The FDA approval of pembrolizumab for adult and pediatric patients with tumor mutational burden (TMB) ≥10: a decision centered on empowering patients and their physicians. Ann. Oncol. 31, 1115–1118 (2020).

95. Sha, D. et al. Tumor Mutational Burden as a Predictive Biomarker in Solid Tumors. Cancer Discov. 10, 1808–1825 (2020).

96. Thorsson, V. et al. The Immune Landscape of Cancer. Immunity 48, 812–830.e14 (2018).

97. Galon, J. & Bruni, D. Approaches to treat immune hot, altered and cold tumours with combination immunotherapies. Nat. Rev. Drug Discov. 18, 197–218 (2019).

98. Rooney, M. S., Shukla, S. A., Wu, C. J., Getz, G. & Hacohen, N. Molecular and genetic properties of tumors associated with local immune cytolytic activity. Cell 160, 48–61 (2015).

99. Johnson, B. J. et al. Single-cell perforin and granzyme expression reveals the anatomical localization of effector CD8^+^ T cells in influenza virus-infected mice. Proc. Natl. Acad. Sci. U. S. A. 100, 2657–2662 (2003).

100. Jayasingam, S. D. et al. Evaluating the Polarization of Tumor-Associated Macrophages Into M1 and M2 Phenotypes in Human Cancer Tissue: Technicalities and Challenges in Routine Clinical Practice. Front. Oncol. 9, 1512 (2019).

101. Pender, A. et al. Genome and Transcriptome Biomarkers of Response to Immune Checkpoint Inhibitors in Advanced Solid Tumors. Clin. Cancer Res. (2020) doi:10.1158/1078-0432.CCR-20-1163.

102. Li, H., van der Merwe, P. A. & Sivakumar, S. Biomarkers of response to PD-1 pathway blockade. Br. J. Cancer 126, 1663–1675 (2022).

103. Fares, C. M., Fenerty, K. E., Chander, C., Theisen, M. K. & Konecny, G. E. Homologous recombination deficiency and molecular subtype are associated with immunogenicity in ovarian cancer. Biomark. Med. 16, 771–782 (2022).

104. Tate, J. G. et al. COSMIC: the Catalogue Of Somatic Mutations In Cancer. Nucleic Acids Res. 47, D941–D947 (2019).

105. Alexandrov, L. B. et al. The repertoire of mutational signatures in human cancer. Nature 578, 94–101 (2020).

106. Nik-Zainal, S. et al. Mutational processes molding the genomes of 21 breast cancers. Cell 149, 979–993 (2012).

107. Ayers, M. et al. IFN-γ-related mRNA profile predicts clinical response to PD-1 blockade. J. Clin. Invest. 127, 2930–2940 (2017).

108. Morad, G., Helmink, B. A., Sharma, P. & Wargo, J. A. Hallmarks of response, resistance, and toxicity to immune checkpoint blockade. Cell 184, 5309–5337 (2021).

109. Newman, A. M. et al. Determining cell type abundance and expression from bulk tissues with digital cytometry. Nat. Biotechnol. 37, 773–782 (2019).

110. Silva-Santos, B., Mensurado, S. & Coffelt, S. B. γδ T cells: pleiotropic immune effectors with therapeutic potential in cancer. Nat. Rev. Cancer 19, 392–404 (2019).

111. Kabelitz, D., Serrano, R., Kouakanou, L., Peters, C. & Kalyan, S. Cancer immunotherapy with γδ T cells: many paths ahead of us. Cell. Mol. Immunol. 17, 925–939 (2020).

112. Bareche, Y. et al. Leveraging big data of immune checkpoint blockade response identifies novel potential targets. Ann. Oncol. 33, 1304–1317 (2022).

113. Ray Chaudhuri, A., et al. Replication fork stability confers chemoresistance in BRCA-deficient cells. Nature 535, 382–387 (2016).

114. De Marco Zompit, M. & Stucki, M. Mechanisms of genome stability maintenance during cell division. DNA Repair 108, 103215 (2021).

115. Li, X. et al. OGT controls mammalian cell viability by regulating the proteasome/mTOR/ mitochondrial axis. Proc. Natl. Acad. Sci. U. S. A. 120, e2218332120 (2023).

116. Donato, E. & Trumpp, A. Targeting the Leukemic stem cell protein machinery by inhibition of mitochondrial pyrimidine synthesis. EMBO molecular medicine vol. 14 e16171 (2022).

117. Gwynne, W. D. et al. Cancer-selective metabolic vulnerabilities in MYC-amplified medulloblastoma. Cancer Cell 40, 1488–1502.e7 (2022).

118. Mullen, N. J. & Singh, P. K. Nucleotide metabolism: a pan-cancer metabolic dependency. Nat. Rev. Cancer 23, 275–294 (2023).

119. Ribas, A. & Wolchok, J. D. Cancer immunotherapy using checkpoint blockade. Science 359, 1350–1355 (2018).

120. You, Z. et al. Homologous recombination repair gene mutations as a predictive biomarker for immunotherapy in patients with advanced melanoma. Front. Immunol. 13, 871756 (2022).

121. Li, Y. et al. SWI/SNF complex gene variations are associated with a higher tumor mutational burden and a better response to immune checkpoint inhibitor treatment: a pan-cancer analysis of next-generation sequencing data corresponding to 4591 cases. Cancer Cell Int. 22, 347 (2022).

122. Zheng, X. et al. SETD2 variation correlates with tumor mutational burden and MSI along with improved response to immunotherapy. BMC Cancer 23, 686 (2023).

123. Wang, G. et al. CRISPR-GEMM Pooled Mutagenic Screening Identifies KMT2D as a Major Modulator of Immune Checkpoint Blockade. Cancer Discov. 10, 1912–1933 (2020).

124. Liu, R. et al. Association of KMT2C/D loss-of-function variants with response to immune checkpoint blockades in colorectal cancer. Cancer Sci. 114, 1229–1239 (2023).

125. Goo, Y.-H. et al. Activating signal cointegrator 2 belongs to a novel steady-state complex that contains a subset of trithorax group proteins. Mol. Cell. Biol. 23, 140–149 (2003).

126. Lintas, C. & Persico, A. M. Unraveling molecular pathways shared by Kabuki and Kabuki-like syndromes. Clin. Genet. 94, 283–295 (2018).

127. Paulussen, A. D. C. et al. MLL2 mutation spectrum in 45 patients with Kabuki syndrome. Hum. Mutat. 32, E2018–25 (2011).

128. Margueron, R. & Reinberg, D. The Polycomb complex PRC2 and its mark in life. Nature 469, 343–349 (2011).

129. Dhar, S. S. et al. Trans-tail regulation of MLL4-catalyzed H3K4 methylation by H4R3 symmetric dimethylation is mediated by a tandem PHD of MLL4. Genes Dev. 26, 2749–2762 (2012).

130. Paull, T. T. 20 Years of Mre11 Biology: No End in Sight. Mol. Cell 71, 419–427 (2018).

131. Mantovani, F., Collavin, L. & Del Sal, G. Mutant p53 as a guardian of the cancer cell. Cell Death Differ. 26, 199–212 (2019).

132. Kim, H. et al. Surrogate reporters for enrichment of cells with nuclease-induced mutations. Nat. Methods 8, 941–943 (2011).

133. Engelen, E. et al. Proteins that bind regulatory regions identified by histone modification chromatin immunoprecipitations and mass spectrometry. Nat. Commun. 6, 7155 (2015).

134. Hughes, C. S. et al. Ultrasensitive proteome analysis using paramagnetic bead technology. Mol. Syst. Biol. 10, 757 (2014).

135. Hughes, C. S. et al. Single-pot, solid-phase-enhanced sample preparation for proteomics experiments. Nat. Protoc. 14, 68–85 (2019).

136. Kong, A. T., Leprevost, F. V., Avtonomov, D. M., Mellacheruvu, D. & Nesvizhskii, A. I. MSFragger: ultrafast and comprehensive peptide identification in mass spectrometry-based proteomics. Nat. Methods 14, 513–520 (2017).

137. da Veiga Leprevost, F., et al. Philosopher: a versatile toolkit for shotgun proteomics data analysis. Nat. Methods 17, 869–870 (2020).

138. Morin, R. D. et al. Frequent mutation of histone-modifying genes in non-Hodgkin lymphoma. Nature 476, 298–303 (2011).

139. Spina, V. et al. The genetics of nodal marginal zone lymphoma. Blood 128, 1362–1373 (2016).

140. Beà, S. et al. Landscape of somatic mutations and clonal evolution in mantle cell lymphoma. Proc. Natl. Acad. Sci. U. S. A. 110, 18250–18255 (2013).

141. Parsons, D. W. et al. The genetic landscape of the childhood cancer medulloblastoma. Science 331, 435–439 (2011).

142. George, J. et al. Comprehensive genomic profiles of small cell lung cancer. Nature 524, 47–53 (2015).

143. DepMap, B. DepMap 22Q2 Public. FigShare https://figshare.com/articles/dataset/DepMap_22Q2_Public/19700056/2 (2022).

144. Meyers, R. M. et al. Computational correction of copy number effect improves specificity of CRISPR-Cas9 essentiality screens in cancer cells. Nat. Genet. 49, 1779–1784 (2017).

145. Nusinow, D. P. et al. Quantitative Proteomics of the Cancer Cell Line Encyclopedia. Cell 180, 387–402.e16 (2020).

146. Ura, H., Togi, S. & Niida, Y. Targeted Double-Stranded cDNA Sequencing-Based Phase Analysis to Identify Compound Heterozygous Mutations and Differential Allelic Expression. Biology 10, (2021).

147. Brown, K. K. et al. Approaches to target tractability assessment - a practical perspective. Medchemcomm 9, 606–613 (2018).

148. Schneider, M. et al. The PROTACtable genome. Nat. Rev. Drug Discov. 20, 789–797 (2021).

149. Li, H. & Durbin, R. Fast and accurate short read alignment with Burrows-Wheeler transform. Bioinformatics 25, 1754–1760 (2009).

150. Cerami, E. et al. The cBio cancer genomics portal: an open platform for exploring multidimensional cancer genomics data. Cancer Discov. 2, 401–404 (2012).

151. Gao, J. et al. Integrative analysis of complex cancer genomics and clinical profiles using the cBioPortal. Sci. Signal. 6, l1 (2013).

152. Middha, S. et al. Reliable Pan-Cancer Microsatellite Instability Assessment by Using Targeted Next-Generation Sequencing Data. JCO Precis Oncol 2017, (2017).

153. Kautto, E. A. et al. Performance evaluation for rapid detection of pan-cancer microsatellite instability with MANTIS. Oncotarget 8, 7452–7463 (2017).

154. Díaz-Gay, M. et al. Assigning mutational signatures to individual samples and individual somatic mutations with SigProfilerAssignment. bioRxiv 2023.07.10.548264 (2023) doi:10.1101/2023.07.10.548264.

155. Goldman, M. J. et al. Visualizing and interpreting cancer genomics data via the Xena platform. Nat. Biotechnol. 38, 675–678 (2020).

156. Pleasance, E. et al. Whole-genome and transcriptome analysis enhances precision cancer treatment options. Ann. Oncol. 33, 939–949 (2022).

157. Kim, S. et al. Strelka2: fast and accurate calling of germline and somatic variants. Nat. Methods 15, 591–594 (2018).

158. Benjamin, D. et al. Calling Somatic SNVs and Indels with Mutect2. bioRxiv 861054 (2019) doi:10.1101/861054.

159. Cleary, J. G., et al. Comparing Variant Call Files for Performance Benchmarking of Next-Generation Sequencing Variant Calling Pipelines. bioRxiv 023754 (2015) doi:10.1101/023754.

160. Titmuss, E. et al. TMBur: a distributable tumor mutation burden approach for whole genome sequencing. BMC Med. Genomics 15, 190 (2022).

161. Davies, H. et al. HRDetect is a predictor of BRCA1 and BRCA2 deficiency based on mutational signatures. Nat. Med. 23, 517–525 (2017).

162. Wu, T. et al. clusterProfiler 4.0: A universal enrichment tool for interpreting omics data. Innovation (N Y) 2, 100141 (2021).

163. R Core Team. R: A Language and Environment for Statistical Computing. (R Foundation for Statistical Computing, 2020).

164. Perez-Riverol, Y. et al. The PRIDE database and related tools and resources in 2019: improving support for quantification data. Nucleic Acids Res. 47, D442–D450 (2019).

165. Freeberg, M. A. et al. The European Genome-phenome Archive in 2021. Nucleic Acids Res. 50, D980–D987 (2022).

